# Striatal Nurr1 Facilitates the Dyskinetic State and Exacerbates Levodopa-Induced Dyskinesia in a Rat Model of Parkinson’s Disease

**DOI:** 10.1101/768374

**Authors:** RC Sellnow, K Steece-Collier, F Altwal, IM Sandoval, JH Kordower, TJ Collier, CE Sortwell, AR West, FP Manfredsson

**Affiliations:** Cell and Molecular Biology Program, Michigan State University, East Lansing, MI; Department of Translational Neuroscience, Michigan State University, Grand Rapids, MI; Rosalind Franklin University, North Chicago, IL; Department of Neurobiology, Barrow Neurological Institute, Phoenix, AZ; Rush University Medical Center, Chicago, IL

**Keywords:** Nurr1, levodopa induced dyskinesia, dopamine, Parkinson’s disease

## Abstract

**Background:** The transcription factor Nurr1 has been identified to be ectopically induced in the striatum of dyskinetic rodents expressing L-DOPA-induced dyskinesia (LID). In the present study, we sought to characterize Nurr1 as a causative factor in LID expression.

**Methods:** We used rAAV2/5 to overexpress Nurr1 or GFP in the parkinsonian striatum of LID-resistant Lewis or LID-prone Fischer-344 (F344) rats. In a second cohort, rats received the Nurr1 agonist amodiaquine (AQ) together with L-DOPA or ropinirole. All rats received a chronic DA agonist and were evaluated for LID severity. Finally, we performed single unit recordings and dendritic spine analyses in drug-naïve rAAV-injected parkinsonian rats.

**Results:** rAAV-GFP injected LID-resistant Lewis rats displayed mild LID and no induction of striatal Nurr1. However, Lewis rats transduced to overexpress Nurr1 developed severe LID. Nurr11 agonism with AQ exacerbated LID in F344 rats. We additionally determined that in L-DOPA-naïve rats striatal rAAV-Nurr1 overexpression 1) increased firing activity in dopamine-depleted striatal direct pathway neurons, and 2) decreased spine density and thin-spine morphology on striatal medium spiny neurons, mimicking changes seen in dyskinetic rats. Finally, we provide post-mortem evidence of Nurr1 expression in the striatum of L-DOPA treated PD patients.

**Conclusions:** Our data demonstrate that ectopic induction of striatal Nurr1 is capable of inducing LID behavior and associated neuropathology, even in resistant subjects. These data support a direct role of Nurr1 in aberrant neuronal plasticity and LID induction, providing a potential novel target for therapeutic development.

## Introduction

Levodopa (L-DOPA) is considered the gold-standard pharmacotherapy for ameliorating motor symptoms in Parkinson’s disease (PD). The precursor to dopamine (DA), L-DOPA alleviates motor symptoms by restoring basal ganglia DAergic tone following the loss of DA from degenerated substantia nigra pars compacta (SNc) neurons (1, 2). Unfortunately, chronic L-DOPA treatment leads the development of drug-induced abnormal involuntary movements (AIMs) in most PD patients (3, 4) known as L-DOPA-induced dyskinesia (LID). These are disruptive hyperkinetic and dystonic movements associated with supra-physiological plasma L-DOPA levels. However, the underlying cause of LID remains largely unknown.

A previous study looking at gene expression differences between the direct and indirect pathway of the basal ganglia in dyskinetic mice showed a marked increase in expression of the transcription factor Nurr1 in direct pathway medium spiny neurons (MSN) (5). The induction of Nurr1 expression in striatal MSN of dyskinetic animals is of particular interest because Nurr1 is not normally expressed in striatal neurons (6). Similarly, we and others have observed that Nurr1 mRNA colocalizes with markers of both the direct and indirect pathway in the rat dyskinetic striatum following dyskinesiogenic L-DOPA dosing, with a higher abundance in the direct pathway (7–9). These findings provide compelling evidence that L-DOPA in the DA depleted striatum appears to induce ectopic Nurr1 expression, however, the role of Nurr1 in LID formation remains unknown.

Nurr1 is an orphan nuclear transcription factor and a member of the NR4A family (10) and its expression is crucial for DAergic neuronal development and long-term function. Indeed, Nurr1 knockout mice are not viable, and Nurr1+/- heterozygotes show DA dysfunction and loss of SNc DA neurons (11–13). Nurr1 expression in the SNc decreases with age in humans, and in fact, multiple Nurr1 isoforms are associated with familial forms of PD (14, 15). Accordingly, Nurr1-based therapeutics are of great interest as potential disease-modifying treatments for PD. However, while Nurr1 in the SN may be of benefit in PD, its activity in the striatum warrants further examination (16–18).

Much research has focused on LID-associated changes in the physiology of the basal ganglia and its connecting target nuclei. *In vivo* recordings of PD patients with deep brain stimulation (DBS) therapy have revealed impaired depotentiation in basal ganglia output centers associated with severe LID (19). Similar findings in preclinical studies demonstrate a loss of bidirectional plasticity associated with LID (20). Specifically, following DAergic denervation, both long-term potentiation (LTP) and long-term depression (LTD) are lost in the striatum, indicating pathological function and abnormal neuroplasticity in corticostriatal transmission. While L-DOPA treatment restores this corticostriatal plasticity, LTD is not present when LID develop (20) and an imbalance or dysfunction of striatal output and is thought to be a key factor in LID expression (21).

Some of these LID-associated changes in striatal plasticity are likely related to changes in dendritic spine morphology. Indeed, striatal spine density and morphology are dramatically altered in animal models following LID induction (22–26). These structural elements are dynamic, and their proliferation or pruning can occur due to normal neuronal processes such as motor learning or can be indicative of pathological states (26–28). As in PD patients (29), animal models of PD have revealed dramatic spine loss following DA depletion, and in preclinical models, reestablishment of spine structure occurs with L-DOPA (22-25, 30,31). However, while L-DOPA administration results in normalization of total spine density in some MSN (23), LID is associated with maladaptive spine changes including a significant increase in the number of large mushroom spines, corticostriatal synapses, and multisynaptic spines in dyskinetic animals (22). Thus, changes in spine density, morphology, and corticostriatal transmission are important pathophysiological factors associated with LID expression. However, our understanding of underlying triggers or causes of these aberrant modifications remains limited.

We posit that Nurr1 is a critical trigger and master regulator necessary for these maladaptive plasticity changes and the induction/expression of LID. Indeed, Nurr1 expression is induced in several cell types by multiple stimuli including stress, addiction, and learning and memory. Within the learning and memory circuits of the hippocampus, Nurr1 is induced in conjunction with associative learning and is required for long-term memory formation (32–34). Nurr1 also plays a key role in the remodeling of basal ganglia circuits during addiction (35). It is thus reasonable to suggest that Nurr1 may play a similar role in the aberrant plasticity associated with dyskinesiogenesis -- a form of maladaptive motor learning. To determine whether Nurr1 is a key contributor to LID development we examined whether it’s overexpression, in the absence or presence of L-DOPA, could induce dyskinesia expression and/or the structural and physiological correlates of LID in striatal MSN. We utilized gene therapy and pharmacological manipulations to modulate expression and activity of Nurr1 in MSN of hemiparkinsonian rats and evaluated the effect on dyskinetic behaviors and MSN structure and function.

## Methods

### Adeno-associated virus production

Vector design and production methods were used as described previously (36). Briefly, the human Nurr1 or the GFP coding sequence were cloned into an rAAV genome under control of the CAG promoter. The genome was packaged into rAAV2/5 via double transfection of HEK293 cells with rAAV genome and the helper plasmid pXYZ5 as previously described (37). Virus was purified using an iodixanol gradient and concentrated in concentration columns. Viral titer was ascertained by dot blot and adjusted to a working titer of 1.0×10^13^ vg/ml.

### Animals and surgeries

Adult male Sprague Dawley (SD), Lewis, and Fischer-344 (F344) rats (200-220g on arrival, Charles River, Wilmington, MA) were used in the studies. Studies were conducted in accordance with Institutional Animal Care and Use Committee (IACUC) approval of Michigan State University (AUF MSU06/16-093-00) and Rosalind Franklin University (AUF# A3279-01). Rats were housed two per cage up until LID behavior testing began, when they were separated and singly housed. Animals were housed in a light-controlled (12 hours light/dark cycle) and temperature-controlled (22±1 °C) room and had free access to standard lab chow and water.

All stereotaxic surgeries were performed under 2% isoflurane. After being anesthetized, animals were placed in a stereotaxic frame and 6-hydroxydopamine (6-OHDA) hydrobromide or rAAV was delivered via a glass capillary needle fitted to a Hamilton syringe (Hamilton, Reno, NV) (38). Three weeks following lesion surgery, animals were tested for spontaneous forepaw use (cylinder test) to estimate lesion efficacy (39). Rats were matched into vector groups based upon cylinder test forepaw deficits in order to ensure equal lesions between the treatment groups. Animals in DA and Nurr1 agonist studies did not receive vector.

Lesions were performed using 5mg/ml 6-OHDA mixed in 0.2mg/ml ascorbic acid immediately preceding injections. SD rats used for spine analysis were lesioned with two 2µL injections of 6-OHDA (10µg per injection) in the left striatum (1^st^ injection from bregma: Anterior Posterior (AP) + 1.6mm, Medial Lateral (ML) 2.4mm, Dorsal Ventral (DV) – 4.2mm from skull; 2^nd^ injection from 1^st^ injection site: AP – 1.4mm, ML +0.2mm, DV – 2.8mm). F344 and Lewis rats used in AIM behavior, LFP, and *in vivo* cell recordings received two 2µl injections of 6-OHDA (10µg per injection), one in the medial forebrain bundle (MFB, from bregma: Anterior Posterior (AP) – 4.3mm, Medial Lateral (ML) + 1.6mm, Dorsal Ventral (DV) - 8.4mm from skull) and one in the SNc (from bregma: AP - 4.8mm, ML + 1.7mm, DV - 8.0mm from skull). The needle was lowered to the site and the injection began after 30 seconds. The needle was removed two minutes after the injection was finished and cleaned between each injection. Lesion efficacy was estimated two and a half weeks following 6-OHDA injection with the cylinder task as previously described (40, 41).

Viral delivery surgeries were performed similarly three weeks following 6-OHDA lesions, as described previously (38). Animals in the LID behavior overexpression studies received a single 2µl injection of either rAAV-Nurr1 or rAAV-GFP targeting lateral striatum (from bregma: AP + 0.5mm, ML + 3.7mm, DV – 5.3mm from skull). Animals used in morphological and electrophysiology studies received two 2µl injections of either rAAV-Nurr1 or rAAV-GFP targeting the entire striatum in order to ensure that all sampled neurons were transduced (1^st^ injection from bregma: AP + 1.0mm, ML + 3.0mm, DV – 4.0mm from dura; 2^nd^ injection from bregma: AP – 1.6mm, ML + 3.8mm, DV – 5mm from dura).

### L-DOPA therapy and abnormal involuntary movements ratings

Animals in electrophysiology and spine analysis experiments were not treated with L-DOPA or DA agonists prior to recordings or sacrifice, and therefore remained non-dyskinetic. For behavioral studies, drug-induced dyskinesia severity was evaluated using the abnormal involuntary movement (AIM) scale (42, 43). F344 and Lewis rats received subcutaneous injections of either increasing doses of L-DOPA (2-8mg/kg) with benserazide (12mg/kg), or the DA receptor agonists SKF-81297 (0.8mg/kg) and quinpirole (0.2mg/kg). Dosing occurred three days a week for three weeks for L-DOPA studies and one week for DA agonist studies. The AIM scale was used to rate drug-induced AIMs, as has been previously described (43, 44). Briefly, AIM severity is evaluated by scoring the level of dystonia of the limbs and body, hyperkinesia of the forelimbs, and hyperoral movements. Each AIM is given two scores—one indicating the intensity (0=absent, 1=mild, 2=moderate, 3=severe) and frequency (0=absent, 1=intermittently present for <50% of the observation period, 2=intermittently present for >50% of the observation period, 3=uninterruptable and present through the entire rating period). Each AIM is given a severity score by multiplying the intensity and frequency, and the overall AIM score for each timepoint is a sum of severity for all behaviors. The sum of all AIM scores from each timepoint makes up the total AIM score. Peak-dose dyskinesia is considered to be 75 minutes post drug administration. An animal is considered non-dyskinetic with a score of ≤4 (44). Animals were observed in 25-minute increments following drug delivery until AIMs subsided.

### Pharmacological activation of Nurr1 with levodopa administration

To assess the impact of pharmacological activation of Nurr1 on LID expression, we utilized the Nurr1 agonist amodiaquine (AQ). Pre-treatment with AQ (20mg/kg twice per day (16)) was initiated prior to introduction of levodopa. The rationale for pretreatment was that potential neuroprotective therapy for PD involving Nurr1 agonists would presumably be initiated soon after diagnosis and prior to introduction of levodopa, which typically occurs approximately one year after diagnosis (45). To determine whether AQ pre-treatment exacerbated LID induction, adult male Sprague Dawley rats rendered unilaterally parkinsonian with 6-OHDA (as described above) first received a low dose (3mg/kg) of levodopa for one week (M-Fr). For all doses, levodopa was administered with 12mg/kg benserazide and 60 minutes after AQ or the vehicle saline. The dose of levodopa was next increased to a moderate dose (6 mg/kg), which was given daily for 2 weeks in the presence of AQ or saline. The dose was finally escalated to a high dose (12mg/kg) for an additional week to determine whether LID severity could be further escalated with high dose levodopa in the presence of AQ. LID were rated at 75 minutes post-levodopa (‘peak dose’) and at timepoints indicated in Figure 3A

### Pharmacological activation of Nurr1 with ropinirole administration

Ropinirole is a non-ergoline D2/D3 dopamine (DA) agonist used for the treatment of early, and as adjunct therapy along with levodopa for advanced PD (46, 47). In early PD, monotherapy with ropinirole has significantly less dyskinesia liability than levodopa in PD patients (46–49) and parkinsonian rodents (50–52). Additional rationale for examining the interaction of ropinirole with Nurr1 agonist neuroprotective therapy is that neuroprotective drugs would most likely be administered to individuals with PD early in the disease, a time when monotherapy with DA agonists like ropinirole is most common.

To determine whether AQ pre-treatment exacerbated dyskinesia induction following chronic ropinirole administration, as above, parkinsonian rats received once-daily AQ or the vehicle saline for 1 week beginning 3-4 weeks after 6-OHDA. To determine whether AQ pre-treatment exacerbated ropinirole-induced dyskinesias (RID) induction, unilaterally parkinsonian rats (as described above) first received a low dose (0.2mg/kg) of ropinirole for three weeks (M-Fr). The dose was escalated to 0.5mg/kg for an additional two weeks. Peak RID occurred at 20 minutes post-injection, and was rated at timepoints indicated in Figure 3D.

### In vivo single unit recordings

Animals used for electrophysiology were shipped to Rosalind Franklin University three weeks following vector delivery and acclimatized for at least 3 weeks prior to recordings. F344 rats were deeply anesthetized with urethane (1.5 g/kg in physiological saline). *In vivo* extracellular single cell recordings of striatonigral projection neurons were measured in vector treated F344 rats without L-DOPA treatment, or in established dyskinetic animals. Electrical stimulation and antidromic activation of striatonigral neurons was performed as previously described (53, 54). For antidromic stimulation, an electrode was placed in the ipsilateral substantia nigra pars reticulata (SNr) (from bregma: AP – 5.0mm, ML + 2.5mm, DV – 8.0mm from dura). For striatal recording ipsilateral to cortical and SNr stimulation, extracellular microelectrodes were lowered slowly through the dorsolateral striatum (from bregma: AP + 0-0.75mm, ML + 3.3-3.7mm, DV – 3.0-6.5mm from dura) while electrical stimuli were delivered to the cortex to isolate responsive striatal single units. Spikes evoked by stimulating the SNr were determined to be antidromically activated based on spike collision with orthodromic spikes occurring consistently over 10 trials (53).

### Tissue collection and immunohistochemistry

Animals received a final injection of either L-DOPA or DA agonists two hours prior to sacrifice. Rats were anesthetized deeply with a lethal dose of sodium pentobarbital, and intracardially perfused with Tyrode’s solution (137mM sodium chloride, 1.8mM calcium chloride dihydrate, 0.32mM sodium phosphate monobasic dihydrate, 5.5mM glucose, 11.9mM sodium bicarbonate, 2.7mM potassium chloride) followed by 4% paraformaldehyde (PFA). Brains were rapidly removed and post-fixed for 72 hours in 4% PFA before being transferred into 30% sucrose. Brains were sectioned on a freezing sliding microtome at 40µm and stored at −20°C in cryoprotectant (30% ethelyne glycol, 0.8mM sucrose in 0.5X tris-buffered saline).

Immunohistochemistry (IHC) was performed as previously reported. A 1:6 series of free-floating tissue was stained for TH (MAB318, MilliporeSigma, Burlington, MA) Nurr1 (AF2156, R&D Systems, Minneapolis, MN) or GFP (AB290, Abcam, Cambridge, United Kingdom). Briefly, sections were washed in 1x TBS with 0.25% Triton x-100, incubated in 0.3% H_2_O_2_ for 30 minutes, and rinsed and blocked in 10% normal goat or donkey serum for 2 hours. Tissue was incubated in primary antibody (TH 1:4000, Nurr1 1.5ug/ml, GFP 1:20,000) overnight at room temperature. After washing, tissue was incubated in secondary antibody (Biotinylated horse anti-mouse IgG 1:500, BA-2001, Vector Laboratories, Burlingame, CA; biotinylated donkey anti-goat IgG 1:500, AP180B, Millipore-Sigma, Burlington, MA; biotinylated goat anti-rabbit IgG 1:500, AP132B, Millipore-Sigma, Burlington, MA) followed by the Vectastain ABC kit (Vector Laboratories, Burlingame, CA). Tissue staining was developed with 0.5 mg/ml 3,3’ diaminobenzidine (DAB, Sigma-Aldrich, St. Louis, MO) and 0.03% H_2_O_2_. Sections were mounted on slides, dehydrated, and coverslipped with Cytoseal (ThermoFisher, Waltham, MA).

### IHC for Nurr1 in human striatal tissue

We examined striatal brain sections from older adults diagnosed with PD or DLB as control. Patient diagnosis was performed by movement disorders specialists at the Rush University Movement Disorders Clinic. All subjects signed an informed consent for clinical evaluation. Postmortem consent was provided by next of kin or a legal representative. The Human Investigation Committee at Rush University Medical Center approved this study.

Free-floating striatal sections were washed in 1X TBS with 0.4% Triton six times for 10 minutes. Sections were incubated 0.3% H_2_O_2_ for 45 minutes, rinsed, mounted on slides, and allowed to dry overnight. Antigen retrieval was performed using Antigen Unmasking Solution (Vector Laboratories) at 80°C for 30 minutes, and then removed from heat and allowed to cool in solution for 30 minutes. Following washing, slides were incubated in 10% NDS serum for 4 hours. Slides were incubated in Nurr1 antibody (1.5ug/ml) overnight. Slides were then washed, incubated in secondary antibody (donkey anti goat 1:500) for 4 hours, followed by incubation in ABC for 1 hour. Nurr1 protein was visualized using Vector SG Peroxidase Substrate Kit (Vector Laboratories). Slides were then coverslipped with Cytoseal (ThermoFisher).

### In situ hybridization

*In situ* hybridization with IHC was performed using the RNAscope® 2.5 HD Duplex Assay according to the manufacturers’ protocol (Advanced Cell Diagnostics, Newark, CA). 40µm striatal sections were treated with the hydrogen peroxide solution for at least 10 minutes, or until active bubbling from the tissue subsided. Sections were washed in 1x TBS, mounted onto slides, and allowed to dry for at least 48 hours. Slides were then boiled for 10 minutes in the Target Retrieval solution, followed by Protease Plus treatment. Tissue was hybridized with target probe for direct and indirect pathway markers (dopamine receptor 1 (D1) or enkephalin (Enk), respectively) (55) for 2 hours at 40°C. Slides were rinsed in 1x RNAscope® Wash Buffer. Sequential amplification steps were then applied to the slides, with 1x Wash Buffer rinses between each amplification. After the sixth amplification, the probe was visualized using the Detect Red Signal solution for 10 minutes.

Immediately following *in situ* hybridization, the tissue was stained immunohistochemically for Nurr1. IHC was performed as described above, and Nurr1 protein was visualized using Vector SG Peroxidase kit (Vector Laboratories, Burlingame, CA).

### Golgi-Cox impregnation and spine analysis

SD rats for spine analysis were perfused as described above with Tyrode’s solution followed by 4% PFA. Brains were removed and hemisected. The caudal portion of the brain was then post-fixed in 4% PFA and used for lesion evaluation. The rostral portion was post-fixed for 1 hour and then transferred to 0.2M phosphate buffe**r** until further processing. The rostral section was sectioned on a vibratome at 100µm. Sections were then processed for Golgi-Cox impregnation as described previously (56). Briefly, sections were sandwiched gently between two glass slides and placed into the Golgi-Cox solution (1% mercury chloride, 1% potassium chromate, 1% potassium dichromate) in the dark for 14 days. Sections were transferred into a 1% potassium dichromate solution for 24 hours. Sections were mounted on 4% gelatin-coated slides and the stain was developed with 28% ammonium hydroxide followed by 15% Kodak fixer. Slides were dehydrated in alcohol and xylene and coverslipped.

Spine density and morphology were quantified using Neurolucida (MicroBrightfield Bioscience, Williston, VT) as has been previously described (22). Neurons of the dorsal striatum were selected for analysis. To be selected for quantification, a neuron needed at least four primary dendrites that projected radially, not bidirectionally, from the cell body. One dendrite per neuron was traced and the spines quantified and typified. Each spine was typified into one of four classes: thin, mushroom, stubby and branched (26). Ten individual dendrites per hemisphere per animal were quantified. Spine quantity and phenotype were evaluated based on total dendrite length, or proximal (dendrite branch orders 1-2) or distal (branch orders 3-n) dendritic regions.

### Stereology

Unbiased stereology was used to determine lesion status via TH loss as previously described (57). Using Stereo Investigator software with the optical fractionator probe (MicroBrightfield Bioscience, Williston, VT), TH-positive neurons in every sixth section of the whole SNc were counted on the intact and lesioned hemisphere, giving an estimate of total TH-positive cells in the SNc. The coefficient of error for each estimate was calculated and was less than 0.1 (Gundersen, m = 1).

### Statistical analysis

Statistical analysis was performed using Statview (version 5.0) or GraphPad Prism version 7.0 (GraphPad Software, La Jolla, CA). All graphs were created in GraphPad. Lesion status was evaluated using unpaired, one-tailed t-tests. Mean AIMs were evaluated using a Mann-Whitney U test or the Kruskal-Wallis test. Differences between vector groups were compared with p≤0.05 being considered statistically significant. For Nurr1 agonist studies, the means and median values at the peak dose LID or RID from each rating day for a given dose of either levodopa or ropinirole were combined for the experimental (AQ+DA agonist) and control (saline+DA agonist) groups to determine of how each group performed as a whole at each dose of DA agonist. Both group means and median values of the peak dose LID or RID were compared for each grouped data set using the non-parametric Mann-Whitney U test. Differences in spine quantity and morphology were evaluated using unpaired t-tests. Differences between vector groups at each recording site and outcomes from animals were compared across experimental groups using a two-way repeated measures ANOVA (GFP vs. Nurr1) x 2 (vehicle vs. drug treatment) with an α set to 0.05 and all “n’s” adequately powered for electrophysiological studies was performed for all experiments. using GraphPad Prism version 7.0 (GraphPad Software, La Jolla, CA), The potential two-way interaction effects were also examined to determine how treatment effects differ as a function of drug treatment or gene therapy (58).

## Results

### rAAV transduction of Nurr1 does not exacerbate AIMs in LID-susceptible but does in LID-resistant rats

We first aimed to determine if ectopic Nurr1 expression can exacerbate LID in LID susceptible and resistant rats. All Fischer-344 (F344) and Lewis rats included in the final analysis were lesioned with ≥94% TH loss (Figure 1B-D), validated post-mortem with stereological quantification of TH immunoreactivity in the SNc. Vector transduction and expression was confirmed with post-mortem IHC in either rAAV-Nurr1 (F344 n=7, Lewis n=5) or rAAV- GFP (F344 n=7, Lewis n=5) transduced rats (Figure 1E-H). Vector-mediated expression results in robust Nurr1 expression in both F344 and Lewis rats. Notably, vector transduction resulted in markedly higher Nurr1 expression than what seen in rAAV-naïve, LID+ F344 animals, where Nurr1 is ectopically upregulated in response to dyskinesiogenic L-DOPA (Supplemental Figure 1A,C). rAAV-naïve Lewis animals, however, express much lower level AIMs than rAAV-naive F344 rats, and also do not ectopically express Nurr1 in the striatum (Supplemental Figure 1B).

**Figure 1.**
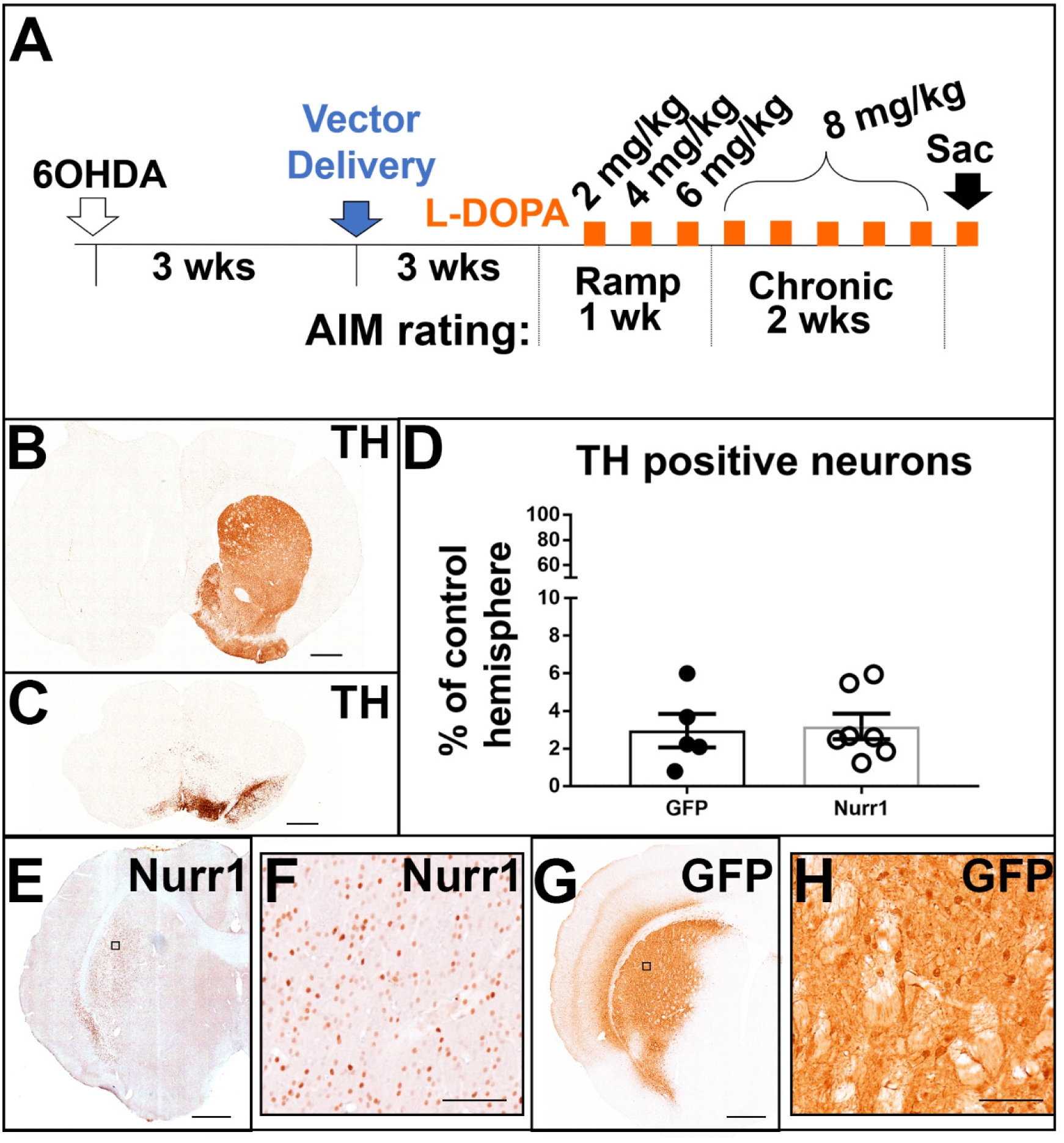
Experimental design and model validation. (A) Experimental timeline showing timing of surgeries and LID induction. AIM scores were evaluated as indicated by arrows. (B-D) 6-OHDA injections caused near-complete loss of TH fibers in the striatum (B) as well as TH-positive cells in the SNc (C). Stereology confirmed that all animals had greater than 94% TH cell loss in the SNc (D). (E-H) Nurr1 (E and F) and GFP (G and H) vector-mediated expression were confirmed in the striatum with IHC. B, C, E, H scale bar=1mm. F, H scale bar=100µm.

AIMs testing began three weeks following virus delivery to ensure maximal transgene expression prior to animals being placed on a L-DOPA treatment paradigm (see Figure 1A for experimental timeline). To evaluate AIMS, animals were first challenged with vehicle (0mg/kg L-DOPA, 12mg/kg benserazide) to determine if Nurr1 overexpression would cause drug-independent AIMs. No vector group showed AIMs following vehicle administration (Supplemental Figure 2). When treated with L-DOPA, F344 treated with rAAV-Nurr1 or GFP rats showed no significant differences in total AIM scores at any rating timepoint (Figure 2A-D) (total AIM sum: D1 2mg/kg rAAV-Nurr1 (Median [Md]=0.5) rAAV-GFP (Md=1), U=17, p>0.05; D3 4mg/kg rAAV-Nurr1 (Md=8) rAAV-GFP (Md=6.5), U=10, p>0.05; D5 6mg/kg rAAV-Nurr1 (Md=39.5) rAAV-GFP (Md=29), U=14, p>0.05; D8 8mg/kg rAAV-Nurr1 (Md=55) rAAV-GFP (Md=42), U=12, p>0.05; D10 8mg/kg rAAV-Nurr1 (Md=61.5) rAAV-GFP (Md=52.5), U=11, p>0.05; D12 8mg/kg rAAV-Nurr1 (Md=57.5) rAAV-GFP (Md=48), U=14, p>0.05; D15 8mg/kg rAAV-Nurr1 (Md=55) rAAV-GFP (Md=63), U=15, p>0.05; D17 8mg/kg rAAV-Nurr1 (Md=62.5) rAAV-GFP (Md=54), U=13, p>0.05). Additionally, Nurr1 overexpression did not differentially impact individual attributes of LID (Supplemental Figure 3). These data suggest that additional induction of striatal Nurr1 above an apparent threshold does not exacerbate AIMs of LID-prone F344 rats.

**Figure 2.**
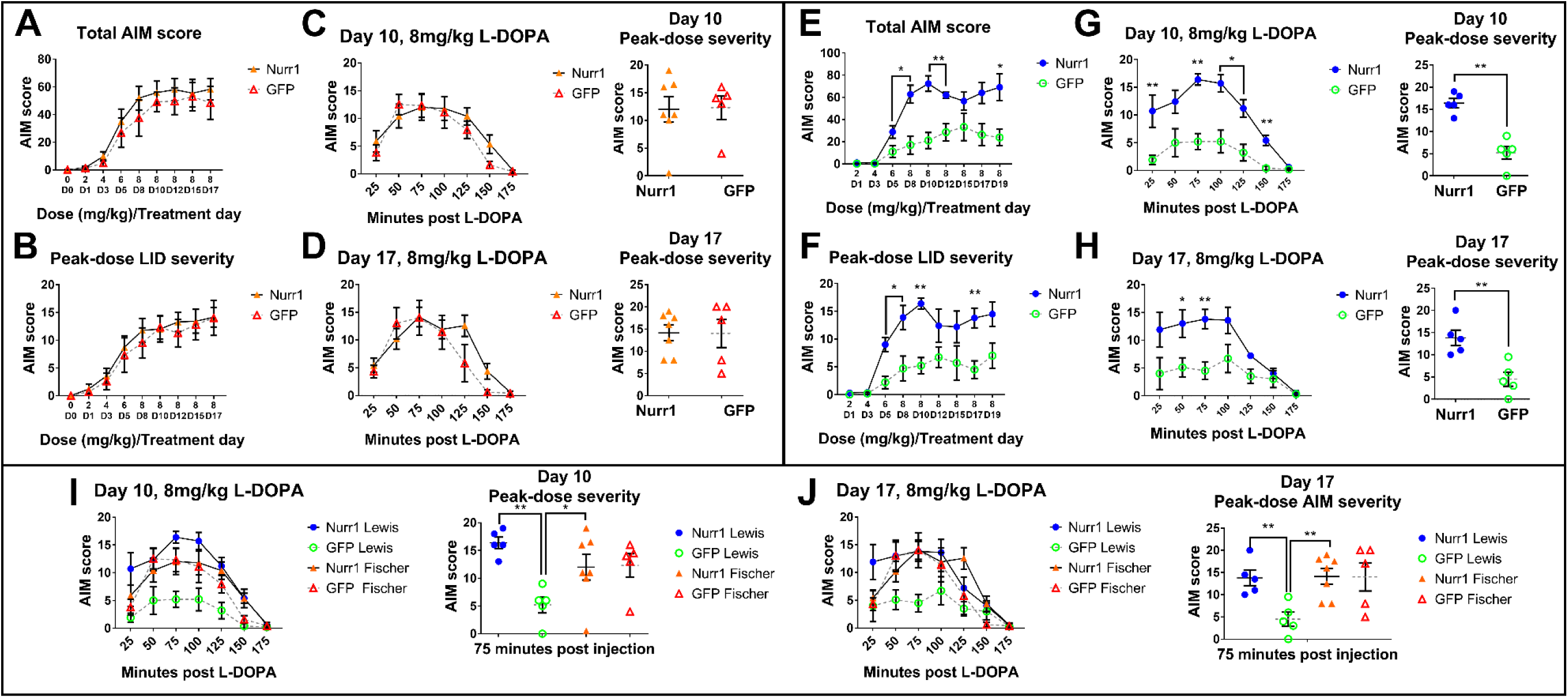
rAAV-mediated striatal overexpression of Nurr1 induces severe AIMs in LID-resistant rats. (A) The total AIM score sum of each rating session showed no differences between rAAV-Nurr1 and rAAV-GFP LID-susceptible F344 rats. Both groups developed severe AIMs similarly over the treatment regimen. (B) Peak-dose LID severity (75 minutes post injection) over each rating period. No differences in peak-dose severity between vector groups was observed in F344 rats. (C-D) Animals were treated with 8mg/kg L-DOPA on days 10-17. Both groups showed similar AIM expression over time (left panels) and peak-dose AIMs (right panels). (E) Total AIM score sum showed exacerbated AIMs on days 5, 8, 10, 12, and 19 in rAAV-Nurr1 treated LID-resistant Lewis rats. (F) Peak-dose LID severity show rAAV-Nurr1 treated Lewis rats developing significantly more severe AIMs than rAAV-GFP animals at 6 and 8mg/kg doses. (C-D) Individual rating periods on days 10-17 with 8mg/kg treatment. rAAV-Nurr1 animals developed AIMs more severe than rAAV-GFP animals at multiple timepoints during the observation period. (I, J) Day 10 (I) and 17 (J) AIM rating time course comparing rAAV treated LID-resistant Lewis rats to LID-susceptible F344 rats. rAAV-Nurr1 Lewis animals showed indistinguishable AIMs time course and peak-dose severity compared to both groups of F344 rats. rAAV-GFP Lewis animals showed significantly lower LID severity than both rAAV-Nurr1 treated Lewis and F344 rats. rAAV-GFP treated F344 rats did not show significantly higher AIMs than rAAV-GFP treated Lewis rats. (*=p≤0.05, **=p≤.01, ***=p≤.001)

In contrast, the normally LID-resistant Lewis rat treated with rAAV-Nurr1 developed severe LID when treated chronically with L-DOPA, whereas rAAV-GFP control rats expressed low-level AIMs (Figure 2E-H). This was first observed at the 6mg/kg treatment, in both total AIM score sum and peak-dose LID (total AIM sum 6mg/kg rAAV-Nurr1 (Md=34) rAAV-GFP (Md=3), U=3, p<0.05; peak-dose AIM 6mg/kg rAAV-Nurr1 (Md=13.5) rAAV-GFP (Md=4), U=0, p<0.01). rAAV-Nurr1 treated Lewis rats also displayed more severe AIMs than their rAAV-GFP counterparts on days 8, 10, and 19 with 8mg/kg L-DOPA (peak dose AIMs: day 8 rAAV-Nurr1 (Md=16) rAAV-GFP (Md=3), U=2, p<0.05; day 10 rAAV-Nurr1 (Md=16) rAAV-GFP (Md=6), U=0, p<0.01; day 17 rAAV-Nurr1 (Md=13.5) rAAV-GFP (Md=4), U=0, p<0.01; day 19 rAAV-Nurr1 (Md=16) rAAV-GFP (Md=3), U=2, p<0.05). Additionally, rAAV-Nurr1 Lewis AIM scores were indistinguishable from both rAAV-GFP and rAAV-Nurr1 F344 rats (Figure 2I-J). Together, this data shows that the expression of Nurr1 is sufficient to overcome resistance to severe LID seen in Lewis animals.

The induction of severe AIMs with ectopic Nurr1 expression in an otherwise LID resistant strain shows that Nurr1 is directly involved in LID development.

### Nurr1 agonist therapy exacerbates LID

Based on the evidence that ectopic striatalNurr1 expression can act as a molecular trigger for LID induction in otherwise resistant Lewis rats, we next sought to determine if pharmacological activation of Nurr1—which is being investigated as a neuroprotective therapy for PD—would also exacerbate LID (17). Unilaterally parkinsonian SD animals were pretreated for 1 week with a neuroprotective dose of the Nurr1 agonist AQ (20mg/kg (16)) followed by L-DOPA+AQ combined administration (Figure 3A). AQ pretreatment, followed by daily administration of L-DOPA+AQ showed evidence of exacerbating LID severity, with a trend at low (3mg/kg levodopa; median severity LD+Sal: 1.1 + 0.33, LD+AQ: 2.5 + 0.36; P=0.05) and high doses (12mg/kg; median severity LD+Sal: 16.9 + 1.44, LD+AQ: 21.4 + 0.99; P=0.100) but reaching statistical significance at the moderate dose tested (6mg/kg; median severity LD+Sal: 13.1 + 0.94, LD+AQ: 19.4 + 0.79; P=0.004) (Figure 3B). Similar results were seen for the mean values of peak dose LID severity (Figure 3C).

**Figure 3.**
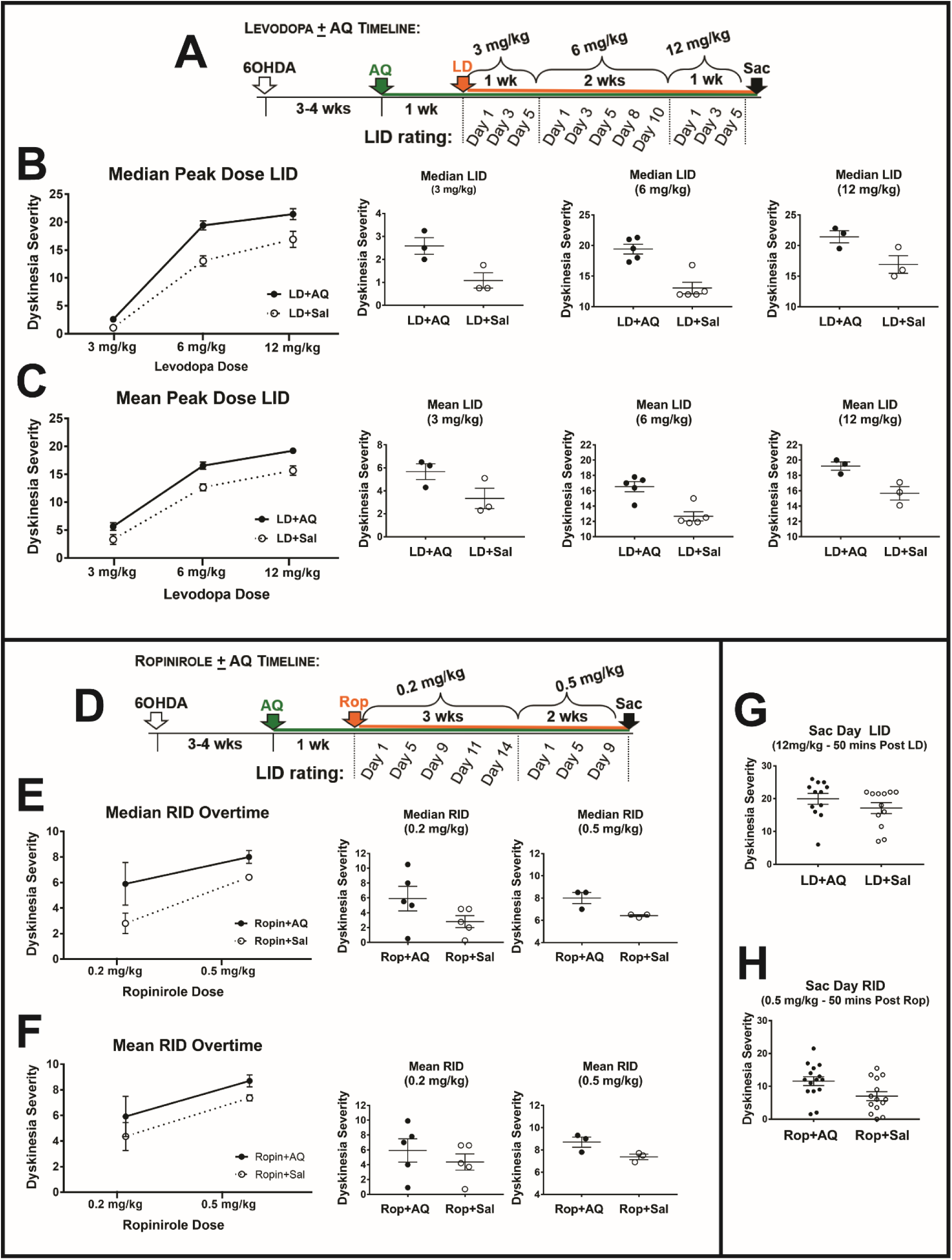
Functional consequences of chronic Nurr1 agonist therapy on the expression of LID and RID in parkinsonian rats. (A, D) Timeline depicting treatment regimen with AQ and L-DOPA (A) or ropinirole (D) in parkinsonian rats. (B-C) Median (B) and mean (C) AIM scores in parkinsonian SD rats trended higher in animals treated with L-DOPA and AQ compared to L-DOPA + saline at low (3mg/kg) and high (12mg/kg) doses of L-DOPA, and were significantly higher at the moderate 6mg/kg dose. (E-F) Median (E) and mean (F) AIM scores in parkinsonian SD rats were significantly higher in rats treated with ropinirole + low-dose AQ compared to ropinirole + saline. LID severity trended higher in animals treated with ropinirole + high-dose AQ compared to ropinirole + saline (G-H) There were no significant differences in AIM scores on the day of sacrifice in animals treated with AQ and L-DOPA (G) or ropinirole (H).

### Nurr1 agonist therapy exacerbates low-dose RID

We next examined the impact of AQ on AIMs in unilaterally parkinsonian SD rats pretreated with AQ (20mg/kg) followed by chronic AQ plus ropinirole, a drug with less dyskinetic liability (Figure 3D). Pretreatment, followed by daily administration of the same neuroprotective dose of the Nurr1 agonist AQ exacerbated RID severity at the low ropinirole dose evaluated (0.2mg/kg; median severity LD+Sal: 2.8 + 0.80, LD+AQ: 5.9 + 1.6; P=0.039) with only a trend of exacerbation at a higher ropinirole dose during chronic treatment (0.5 mg/kg; median severity L-DOPA+Sal: 6.4 + 0.08, L-DOPA+AQ: 8.0 + 0.5; P=0.05). In contrast to median RID data, which is less affected by outliers, there were no significant difference in mean RID values except at the final rating on day of sacrifice (Figure 3H; P=0.018).

### Nurr1 expression is induced by pharmacological activation of direct pathway MSN

We next examined whether preferential activation of either the striatal direct or indirect pathways is sufficient to reproduce the pathophysiological characteristics of LID and upregulation of Nurr1. Previous reports have found L-DOPA-induced Nurr1 expression in both direct and indirect pathway MSN (dMSN and iMSN, respectively). To examine this, we induced AIMs in parkinsonian F344 rats with the selective D1/D5 agonist SKF-81297 or D/D32 agonist quinpirole; saline injection as a control. Rats were treated for one week with either drug and euthanized 2 hours following the final dosing. Moderate and severe AIMs developed in animals treated with SKF-81297 (Figure 4). Quinpirole-treated rats expressed significantly less severe AIMs. No saline-treated animals developed AIMs. Statistical analysis showed that drug-induced AIMs were significantly different between treatment groups (peak-dose AIMs: Kruskal-Wallis statistic=6.78, p<0.05). IHC revealed abundant Nurr1 expression in the lesioned striatum of SKF-81297 rats, but notably not in those expressing low-level AIMS following D2/D3 agonist treatment or in the absence of AIMs. These data suggest that selective D1 receptor activation readily elevates Nurr1 and AIMs behavior, supporting the view that selective indirect pathway activation does not induce maladaptive striatal Nurr1 expression, but that direct pathway activation is required for this event.

**Figure 4.**
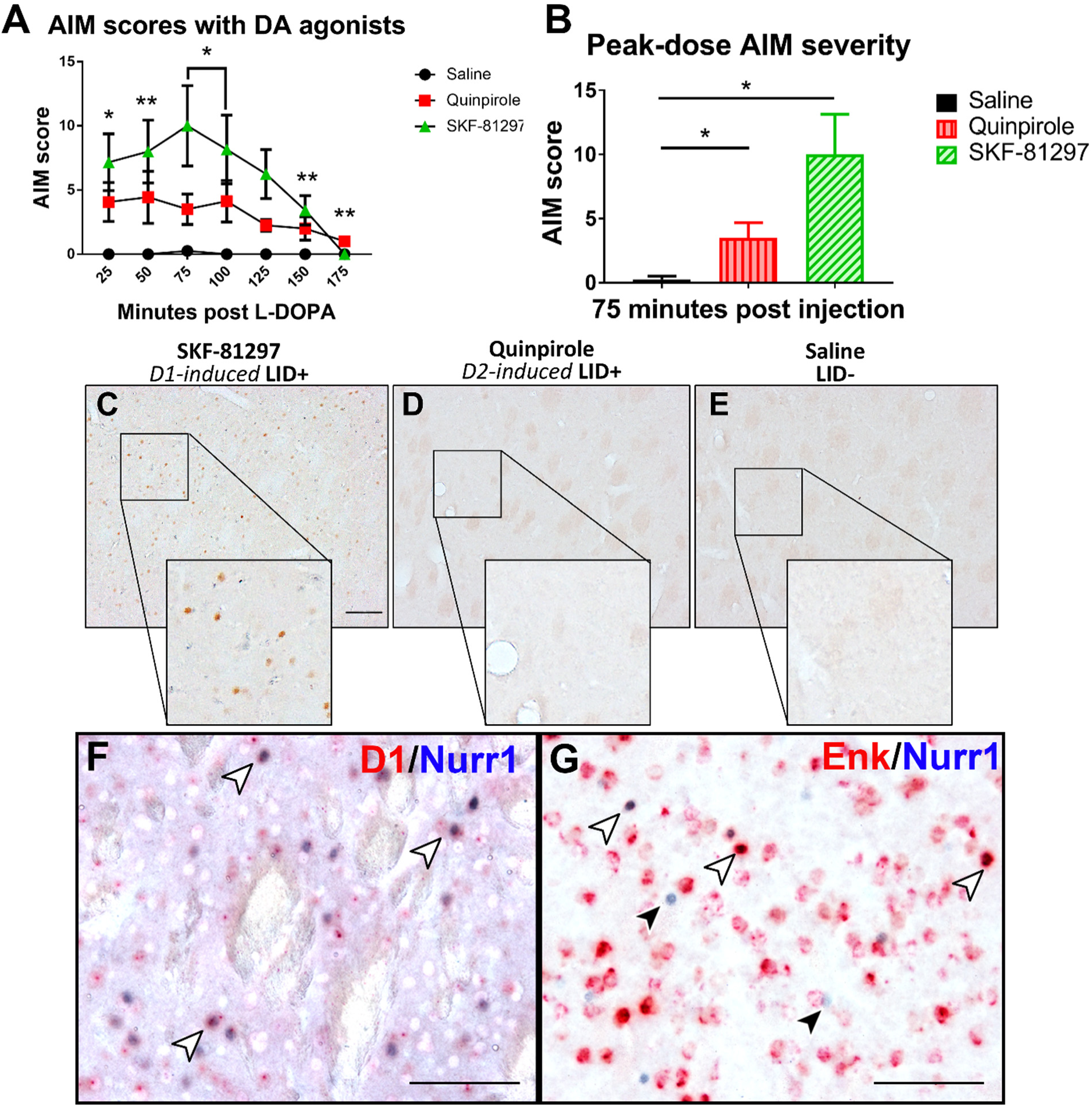
Abnormal striatal Nurr1 expression is induced by direct D1 receptor activation. (A) AIM scores from the final treatment with either D1 agonist (SKF-81297), D2 agonist (quinpirole), or saline. D1-agonist treated rats displayed severe AIMs, while D2 agonist treated animals expressed moderate AIMs. No AIMs were observed in saline treated animals. (B) Peak-dose AIM severity from the final drug treatment (*=p≤0.01, **=p≤.05). (C-E) Nurr1 IHC in lesioned striatum of animals treated with SKF-81297 (C), quinpirole (D) or saline (E). Nurr1 protein was only observed in the lesioned striatum of dyskinetic rats treated with SKF-81297 (C). Nurr1 was not observed in the lesioned striatum of LID+ rats treated with quinpirole (D). No Nurr1 was seen in saline treated animals (E). Scale bar=100µm. (F) *In situ* hybridization for D1 (red) and IHC for Nurr1 protein (blue) in striatum of LID+ animal treated with D1 agonist SKF-81297. D1 transcript and Nurr1 protein are colocalized (white arrows) in some neurons, but not others (black arrows). (G) *In situ* hybridization for enkephalin (red) and IHC for Nurr1 protein (blue) in the striatum of an LID+ animal treated with D1 agonist SKF-81297. The enkephalin transcript is seen to colocalize with Nurr1 protein (white arrows). Some cells show Nurr1 expression with no enkephalin transcript (black arrows) Scale bar=100µm.

To determine if selective direct pathway activation leads to Nurr1 expression only in D1 MSN, we performed dual label *in situ* hybridization with IHC to localize Nurr1 protein with mRNA of direct and indirect pathway markers (dopamine receptor 1 (D1 and enkephalin (Enk), respectively) (55). We observed cellular colocalization of Nurr1 protein and both markers of striatal projection neurons (D1 and Enk) in SKF-81297 AIM-expressing animals (Figure 4F-G). These findings suggest that Nurr1 expression in the indirect pathway is dependent on direct pathway activation. We also examined this in rats treated with quinpirole and saline, but no colocalization was observed, as no Nurr1 protein was expressed in these animals (Supplemental Figure 4).

### rAAV-induced Nurr1 expression in the absence of L-DOPA induces LID-like pathophysiological corticostriatal signaling and dendritic spine changes

Animals for electrophysiology and spine analysis studies were rendered parkinsonian with 6-OHDA injections as described previously (59, 60). Animals included in these analyses showed ∼75-80% cell loss in the injected hemisphere and there was no difference in lesion severity between vector groups (Figure 5A-C) (rAAV-Nurr1 % TH neurons remaining=21.48±3.12; rAAV-GFP % TH neurons remaining=27.87±5.26; t_13)_=1.08, p>0.05). This degree of lesion is sufficient to induce enduring striatal changes that occur following DA depletion. Transgene expression was confirmed with IHC, and all animals included in the analysis showed robust transgene expression in the striatum (Figure 5D-E). These animals were used for *in vivo* electrophysiology (rAAV-Nurr1 n=15, rAAV-GFP n=9) or spine analysis (rAAV-Nurr1 n=4, rAAV-GFP n=3).

**Figure 5.**
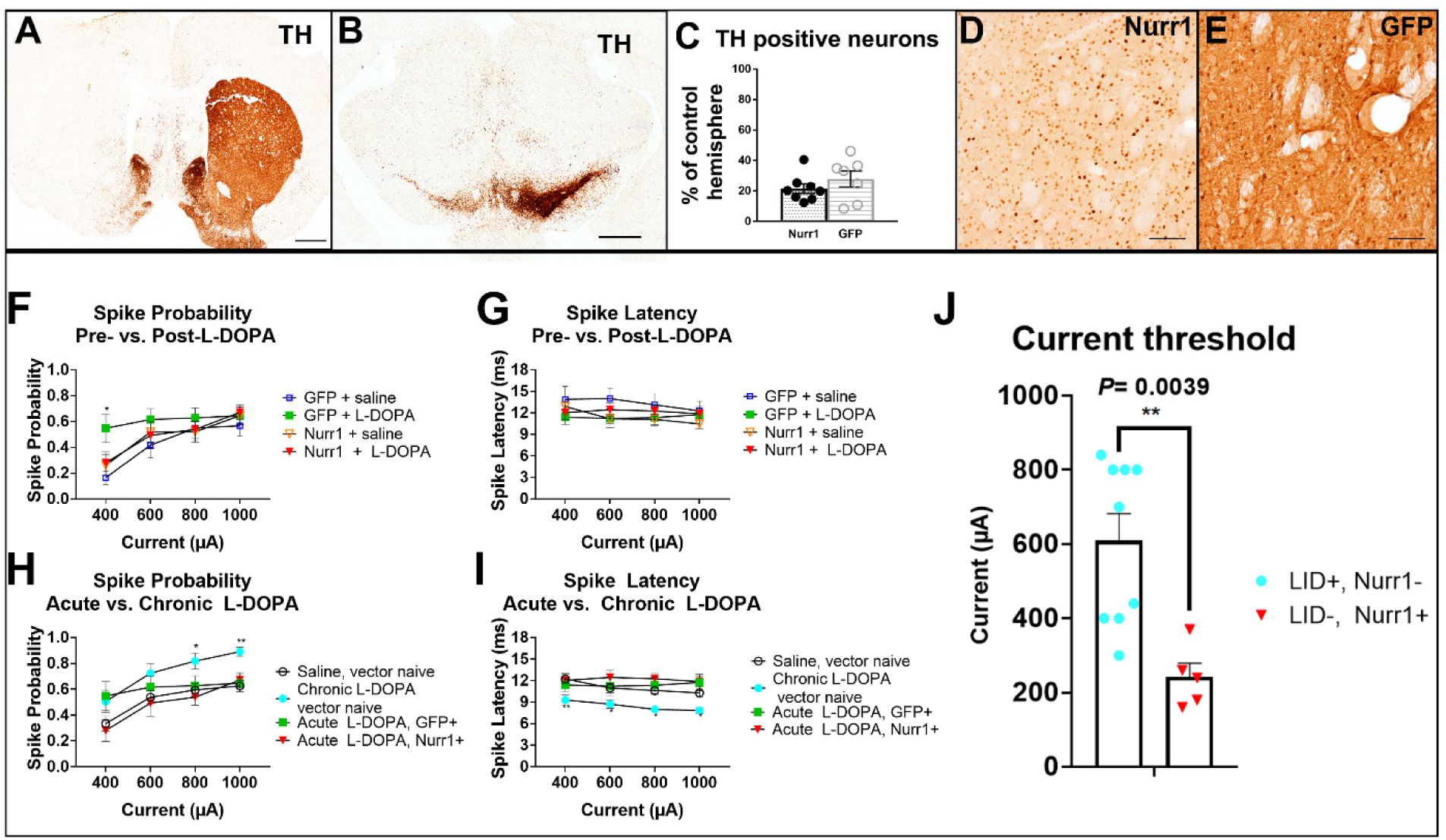
Cortically-evoked response of striatonigral MSN to antidromic stimulation. (A-C) Lesion status was confirmed with IHC for TH. TH immunoreactivity was dramatically reduced in the interjected hemisphere of the striatum (A) and substantia nigra (B). Stereological estimates of remaining TH-positive neurons show significant cell loss in both vector groups (C). (D-E) Transgene expression from viral vector delivery was confirmed in the striatum with IHC for Nurr1 (D) or GFP (E). A and B scale bar=1mm. D and E scale bar=1µm. (F-J) Comparisons between cortically-evoked spike characteristics of antidromically-activated striatonigral projection neurons recorded from DA-depleted parkinsonian, vector naïve rats treated with saline or chronic L-DOPA, and vector-treated (rAAV-GFP or rAAV-Nurr1) animals not chronically treated with L-DOPA. (F) Spike probability pre- and post-L-DOPA delivery in GFP and Nurr1 injected rats. GFP rats showed a higher spike probability after L-DOPA delivery at 400µA stimulation. (G) No differences in spike latency before and after L-DOPA injection were observed in either vector group. (H) Established LID+ rats (Chronic L-DOPA, vector naïve) showed higher spike probability than GFP injected, Nurr1 injected, and rAAV-naïve saline treated controls. (I) LID+ rats were showed significantly lower spike latency than both vector-treated groups and saline control (*=p≤0.05, **=p≤.01). (J) The current threshold of rAAV-Nurr1 rats (242µA ± 36.9) was significantly lower than that of vector naïve, LID+ rats (608.9µA ± 72.9, p=0.0039).

To understand how ectopic Nurr1 expression can impact MSN activity and signaling, we performed *in vivo* extracellular single until recordings of unidentified MSN in L-DOPA naïve parkinsonian rats injected with rAAV-Nurr1 or rAAV-GFP. After a cell was isolated, cortically-evoked responses to a range of stimulation intensities (1000µA, 800µA, 600µA, and 400µA) were recorded. We compared the spike probability and onset latency of cortically-evoked spikes in both groups before and after delivering acute low dose (5mg/kg) L-DOPA (Figure 5-G). There was no difference in either spike probability or latency between rAAV-GFP and rAAV-Nurr1 rats. However, compared to rAAV-GFP animals, rAAV-Nurr1 rats exhibited decreased spike probability with a stimulation intensity of 400µA following L-DOPA delivery (Figure 8F). We were able to positively identify a small number of D1 MSN. In this population, rAAV-Nurr1 cortically evoked spike response mimicked the spike profile of rAAV-naïve dyskinetic rats (Supplemental Figure 5).

We next compared vector treated rats with vector naïve animals treated chronically with L-DOPA, and therefore LID+. Spike probability was increased, and spike onset latency was decreased in LID+ animals, indicating an increase in corticostriatal drive. Both spike probability and latency in the rAAV-GFP and rAAV-Nur1 animals were significantly different than the firing patterns of the LID+ animals, and more resembled vector naïve, L-DOPA naïve rats (Figure 5H-I). This suggests that pathophysiological alterations to corticostriatal circuits affecting striatal projection neuron activity are dependent on LID presence, and not Nurr1 expression alone.

Finally, we measured the current threshold of identified striatonigral MSN (dMSN) in vector naïve, LID+ rats and compared them to similar measures from non-dyskinetic rats treated with rAAV-Nurr1. We found that in the striatum of rAAV-Nurr1 rats there was a significantly reduced current threshold required to elicit antidromic spike activity in dMSN as compared to animals with established dyskinesias (rAAV-Nurr1 L-DOPA naïve, 242µA ± 36.9; rAAV-naïve LID+, 608.9µA ± 72.9; p=0.0039; Figure 5J). This indicates that dMSN ectopically expressing Nurr1 are significantly more excitable than those in a LID+ rat, suggesting Nurr1 may be priming the abnormal signaling activity that occurs with chronic L-DOPA therapy associated with LID.

### MSN spine density and morphology changes are induced by Nurr1 expression

Dramatic changes in spine density and morphology in LID models has been previously demonstrated (22, 61). As Nurr1 is involved in learning and memory-associated plasticity classically associated with spine changes (32, 34), we sought to determine if Nurr1 expression could affect MSN spines without exposure to L-DOPA. Similar to the rats used for electrophysiology, these animals did not receive L-DOPA at any time, again allowing us to determine the effect of Nurr1 on DA-depleted MSN independent of drug treatment. Spine analysis revealed there was a significant decrease in total spine density in rAAV-Nurr1 treated animals (rAAV-Nurr1 total spines/10µm=4.85±0.10, rAAV-GFP total spines/10µm=5.98±0.31; t(_5_)=3.68, p<0.05) (Figure 6A). This difference was not specific to either the proximal or distal portions of the dendrite (rAAV-Nurr1 proximal spines/10µm=3.54±0.31, distal spines/10µm=5.58±0.41; rAAV-GFP proximal spines/10µm=4.24±1.45, distal spines/10µm=6.89±0.74). These data clearly indicate that Nurr1 dose impact dendritic spine dynamics, even in the absence of DA signaling related to L-DOPA, and may play a role in spine plasticity changes accompanying LID.

**Figure 6.**
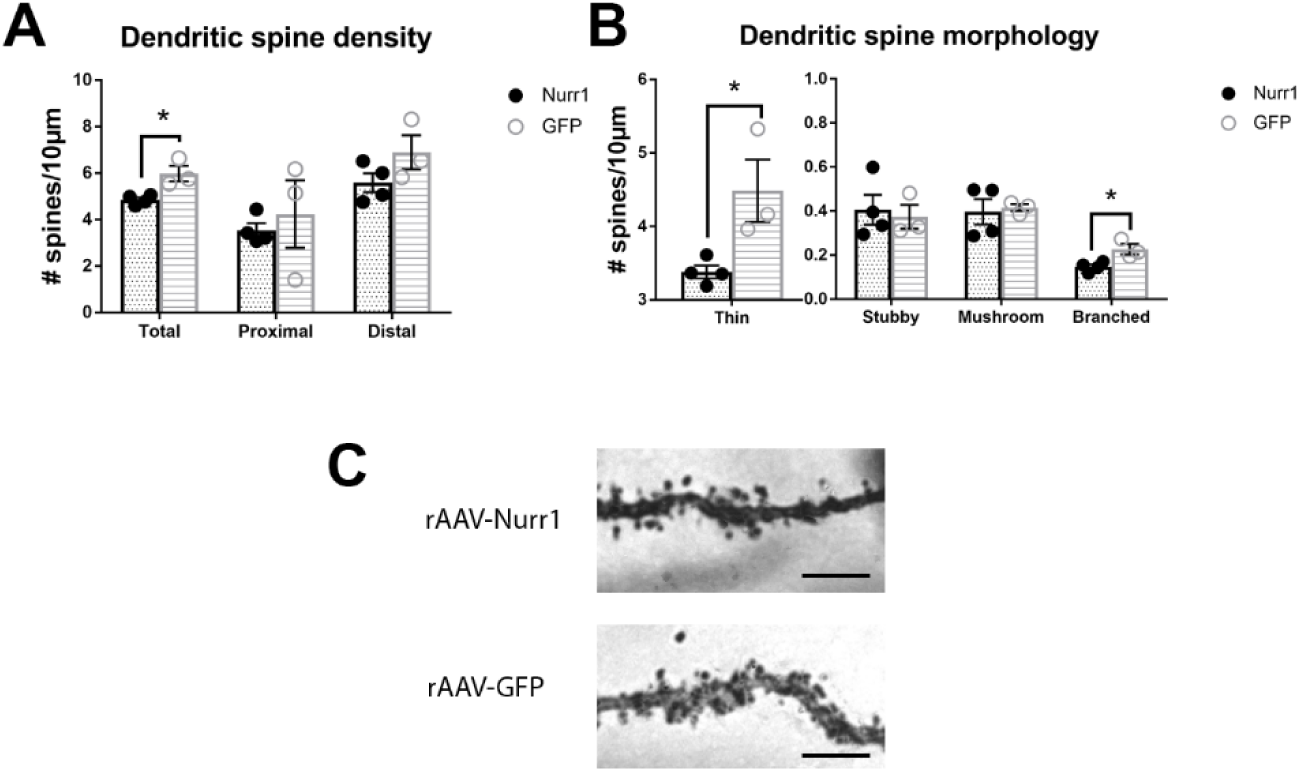
Nurr1-induced alterations in dendritic spine density and morphology. (A) Dendritic spine density of MSN in the lesioned hemisphere of rats treated with rAAV-Nurr1 or rAAV-GFP. Nurr1 expression caused a decrease in total number of spines. rAAV-Nurr1 total spines/10µm=4.85±0.10, rAAV-GFP total spines/10µm=5.98±0.31; t_(5)_=3.68, *=p≤0.05 (B) rAAV-Nurr1 expression induced a decrease in both thin and branched type spines compared to rAAV-GFP controls. rAAV-Nurr1 thin spines/10µm=3.38±0.09, rAAV-GFP thin spines/10µm=4.48±0.43; t_(5)_=2.98, *=p≤0.05. (C) Representative image of dendritic spines on Golgi-Cox stained MSN. Scale bar=50µm.

We next compared spine morphology, an indicator of synaptic strength and spine dynamics (26), between rAAV-Nurr1 and rAAV-GFP animals. We found that ectopic Nurr1 expression lead to a change in two specific spine phenotypes. rAAV-Nurr1 animals displayed significantly fewer thin spines than controls (rAAV-Nurr1 thin spines/10µm=3.38±0.09, rAAV-GFP thin spines/10µm=4.48±0.43; t_(5)_=2.98, p<0.05). We also observed fewer branched (also known as cupped or bifurcated) spines in rAAV-Nurr1 (branched spines/10µm=0.15±0.01) than in rAAV-GFP animals (branched spines/10µm=0.23±0.02; t_(5)_=3.35, p<0.05) (Figure 6B). No difference between groups was observed with stubby or mushroom spines. Cumulatively, these data implicate Nurr1, independent of L-DOPA, as a molecular regulator of the maladaptive spine plasticity that has been shown in animal models of LID.

### Nurr1 is expressed in the striatum of L-DOPA-treated PD patients

Conservation of a variable across species is important to support not only its involvement in preclinical models but its relevance to the human disease state. To assess this, we used IHC to determine if Nurr1 expression is present in the striatum of dyskinetic PD patients who had been treated with L-DOPA. In multiple cases (all confirmed dyskinetic), we observed distinct nuclear Nurr1 staining in the putamen of PD patients (Figure 7A). This signal was not observed in an age-matched controls with dementia with Lewy bodies (DLB) that never received L-DOPA (Figure 7B). This observation is the first direct connection of Nurr1 to LID in PD patients.

**Figure 7.**
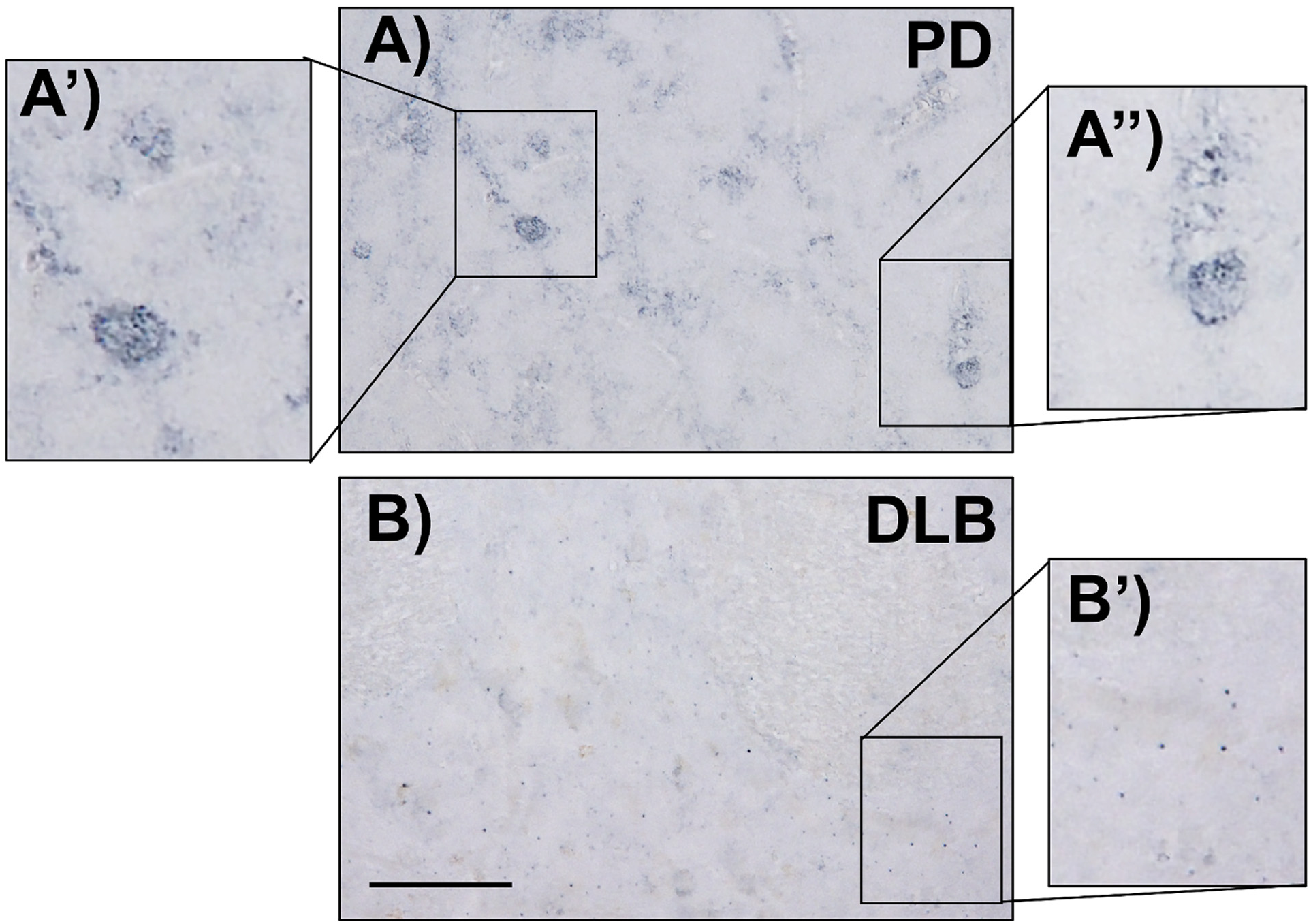
Nurr1 is expressed in the striatum of L-DOPA treated PD patients. IHC for Nurr1 in the putamen of post-mortem patient samples of two Parkinson’s disease patients (A-B) and an age-matched patient with dementia with Lewy body (DLB) (L-DOPA naïve) (C). White arrows indicate nuclear staining for Nurr1 in the putamen of both PD patients. Scale bar=50µm.

## Discussion

While previous studies have demonstrated that ectopic induction of striatal Nurr1 is associated with LID (5, 9), the current study demonstrates that not only is striatal Nurr1 expression capable of inducing LID behavior in genetically resistant subjects, but also provides the first evidence of a genetic modifier of LID directly inducing morphological and electrophysiological signatures of this behavioral malady independent of L-DOPA. In the following paragraphs we provide discussion related to the contribution of our studies in advancing understanding of the role of this transcription factor in LID, as well as potential clinical ramification.

### Genotype-to-Phenotype Impact of Ectopic Striatal Nurr1

The homogenous genotype of inbred rat strains has served in preclinical models to advance understanding genetic factors underlying differences in behavioral phenotypes. Lewis and F344 rats are well known to display differences in response to psychostimulant addiction liability (62–65), a behavioral phenomenon with many similarities to LID in that both are considered to result from aberrant associative or motor learning and rewiring of basal ganglia neural circuits following DA-related priming. We recently demonstrated (Steece-Collier et al., manuscript submitted) that these rat strains also show dramatically different LID liability, with Lewis rats showing distinct resistance to LID expression in contrast to robust development in F344 rats. Using this novel model, we report here that ectopic induction of striatal Nurr1 in LID-resistant Lewis rats can induce LID severity to the same high level as seen in LID-prone F344 rats, suggesting that striatal Nurr1 expression is a key factor in: 1) LID development, and 2) the differential susceptibility between these two inbred rat strains. Further, these data suggest that Nurr1 expression is not a downstream byproduct of LID, but rather plays a causative role in LID development. Continuing studies aimed at identifying genetic or epigenetic differences between the strains that may explain their differential upregulation of Nurr1 in response to L-DOPA would be valuable for the identification of molecular traits or signatures involved in LID susceptibility.

In addition to ectopic induction of striatal Nurr1 being capable of converting genetically LID-resistant rats to LID-expressing rats, the current data provides novel insight that this particular genetic modifier of LID can directly induce phenotypic morphological and electrophysiological changes independent of L-DOPA. Specifically, we found that ectopic Nurr1 expression in the absence of L-DOPA resulted in a decrease in total number of dendritic spines, reflective of specific decreases in thin and branched spines. While a decrease in total spine number may appear contrary to previous reports showing an overall *restoration* in the striatal spine density in dyskinetic animals, at least two studies have reported that despite restoration of spines with LID, the total density remained below control levels (22, 23). Further, spine density on dMSN neurons has been observed to show a specific decrease in dyskinetic mice, additive to that associated with DA depletion (23). An additional study demonstrated a significant decrease in average spine density in LID expressing rats in both dMSN and iMSN (24). While we were unable to differentiate spine changes on specific populations of MSN, the current findings are in keeping with these previous reports and implicate Nurr1 as a molecular factor involved in promoting these phenotypic maladaptive changes.

While the mechanism of this effect remains to be determined, Nurr1 is known to heterodimerize with another NR4A transcription factor, Nur77 (66). Nur77 is endogenously expressed in the adult striatum, and is increased by L-DOPA treatment in denervated animals, primarily in dMSN (67, 68). Similar to our findings, viral overexpression of Nur77 in the striatum can exacerbate AIMs in rats (69) and also induce spine loss in hippocampal neurons, supporting an important role the NR4A family of transcription factors in affecting synaptic plasticity (70).

In keeping with ectopic Nurr1 directly affecting striatal synaptic plasticity we show here that Nurr1 expression alone, independent of dyskinesiogenic L-DOPA, can induce an LID-like alteration in dMSN firing patterns in response to acute, non-dyskinesiogenic, low-dose L-DOPA given at the time of recording. Specifically, these studies provide unique insight that Nurr1 is capable of inducing hyper-cortical drive. Indeed, in a small subset of identified dMSN, we saw an identical increase, as compared to rAAV-GFP treated subjects, in spike probability in rAAV-Nurr1 as in the LID+ rats treated chronically with L-DOPA suggesting that Nurr1 induces LFP-like signaling in D1 neurons, regardless of chronic L-DOPA administration and LID expression.

### Direct pathway activation is required for Nurr1 induction

While there is doubtless interaction of the direct and indirect pathways in LID pathology, Nurr1 had previously been reported to be expressed primarily in MSN of the direct pathway in association with LID (5). Our pharmacological studies employing the selective D1/D5 agonist SKF-81297 or D2/D3 agonist quinpirole support that while AIMs are observed under conditions of either preferential direct or indirect pathway activation, AIMs are more severe with a selective D1 receptor agonist and Nurr1 induction in striatal MSN occurred only in parkinsonian rats treated with the DA D1 receptor agonist. Interestingly, despite preferential activation of D1 receptor MSN, we observed Nurr1 induction in both the direct and indirect pathway MSN. It is uncertain why Nurr1 transcript was not induced following treatment with the selective D2 receptor agonist. It is possible that there are D2-dependent, Nurr1-independent, pathways which can lead to LID development, suggesting that the mechanism for the low-severity LID development in the Lewis strain is a distinct phenotype from the canonical parkinsonian LID. However, while AIMs were observed with quinpirole, the phenotype was not only less severe, but was also generally more reflective of hyperkinesia than the trunk and limb dystonia typical of SKF-81297 or L-DOPA.

These findings indicate that Nurr1 induction in the striatum is dependent on stimulation of striatonigral dMSN, and that D1 receptor activation is sufficient to induce Nurr1 expression in not only direct, but also indirect pathway MSN. These data align with studies of other immediate early genes in LID, such as Homer-1a and FosB, which show that D1 priming is required for expression in D2 neurons (71–73). Based on the finding that F344 rats treated with the D2 selective agonist quinpirole and Lewis rats treated with L-DOPA develop only mild-to-moderate AIMs, which were not associated with striatal Nurr1 induction, it is reasonable to suggests that Nurr1 may act as a ‘master regulator’ in the striatum and when expressed/activated can induce changes in corticostriatal transmission, striatal output, and motor behavior that lead to more severe dyskinesia.

### Clinical Indications for Nurr1 in PD: Nurr1 Agonists and LID

Nurr1 has received a great deal of attention in the past several of years for its essential role in the survival of nigral DA neurons (11–13), and its potential to slow death of these neurons in PD (16-18, 74-76). However, while Nurr1 agonist therapy directed at *nigral* neurons may offer a promising new avenue of neuroprotective therapy, the evidence presented here and in other reports (5, 9) suggests that induction of extranigral Nurr1, specifically within the striatum, is a critical trigger for the induction of LID. We hypothesized that if systemic administration of drugs that activate Nurr1 were to be employed in PD for neuroprotection, this could result in the exacerbation of LID, further compromising quality of life. To provide the first proof-of-principle evidence, we employed the FDA approved antimalarial drug amodiaquine (AQ) that has been demonstrated in human neuroblastoma cells expressing Nurr1 to facilitate the transactivation of Nurr1 thru direct binding to the Nurr1 ligand binding domain. AQ provide neuroprotection against 6-OHDA-induced nigral DA neuron loss in an *in vivo* rat model (16). We demonstrate here that daily AQ given in the presence of either L-DOPA or the D2/3 receptor agonist ropinirole, a drug with less dyskinesia liability and is often used in early PD (refs) significantly exacerbates expression of AIMs in parkinsonian rats. These data suggest that systemic administration of Nurr1 activating drugs with DA agonist therapy may be contraindicated in PD.

### Clinical Indications for Nurr1 in PD: Suppressing Activation of a Repressed Gene

A second clinical facet of Nurr1 activation that holds promise for therapeutic development is related to its apparent repressed status in the normal adult *striatum* and its ectopic induction in the dyskinetic striatum. Specifically, while other NR4A transcription factors are expressed endogenously in the striatum, in the non-dyskinetic striatum Nurr1 is undetectable (6, 9, 77). This unique gene profile could provide a powerful therapeutic advantage since targeted inhibition of striatal Nurr1 would be anticipated to result in no adverse physiological events since its expression within the striatum is aberrant. This would contrast targeted inhibition of other striatal LID-associated genes (e.g.: FosB) that are known to be integral and necessary for normal neuronal function.

### Conclusions

Data presented here demonstrate that Nurr1 is not only a marker of LID in preclinical models and in dyskinetic human PD patients, but is directly involved in LID development, expression, and severity. We further demonstrate that Nurr1 is capable of effecting corticostriatal transmission and striatal projection neuron activity, as well as neuronal spine morphology in a way that mimics and could promote LID. This characterization of Nurr1 highlights its important and potentially, causative role in LID, suggesting that Nurr1 agonists-based therapies for neuroprotection in PD may be contraindicated but that targeted silencing may be uniquely poised to provide a safe and effective anti-dyskinetic treatment option.

## Acknowledgements

The authors would like to acknowledge Kuei Tseng for intellectual input and assistance with. electrophysiological studies. Nathan Kuhn, Brian Daley, Jennifer Stancati, and Christopher Kelp for surgical and histological assistance. This work was supported by the National Institute of Neurological Disorders and Stroke NS098079 (FPM, KSC, ARW) and a Target Advancement Program grant from the Michael J. Fox Foundation (KSC).

---

**Supplemental Figure 1.**
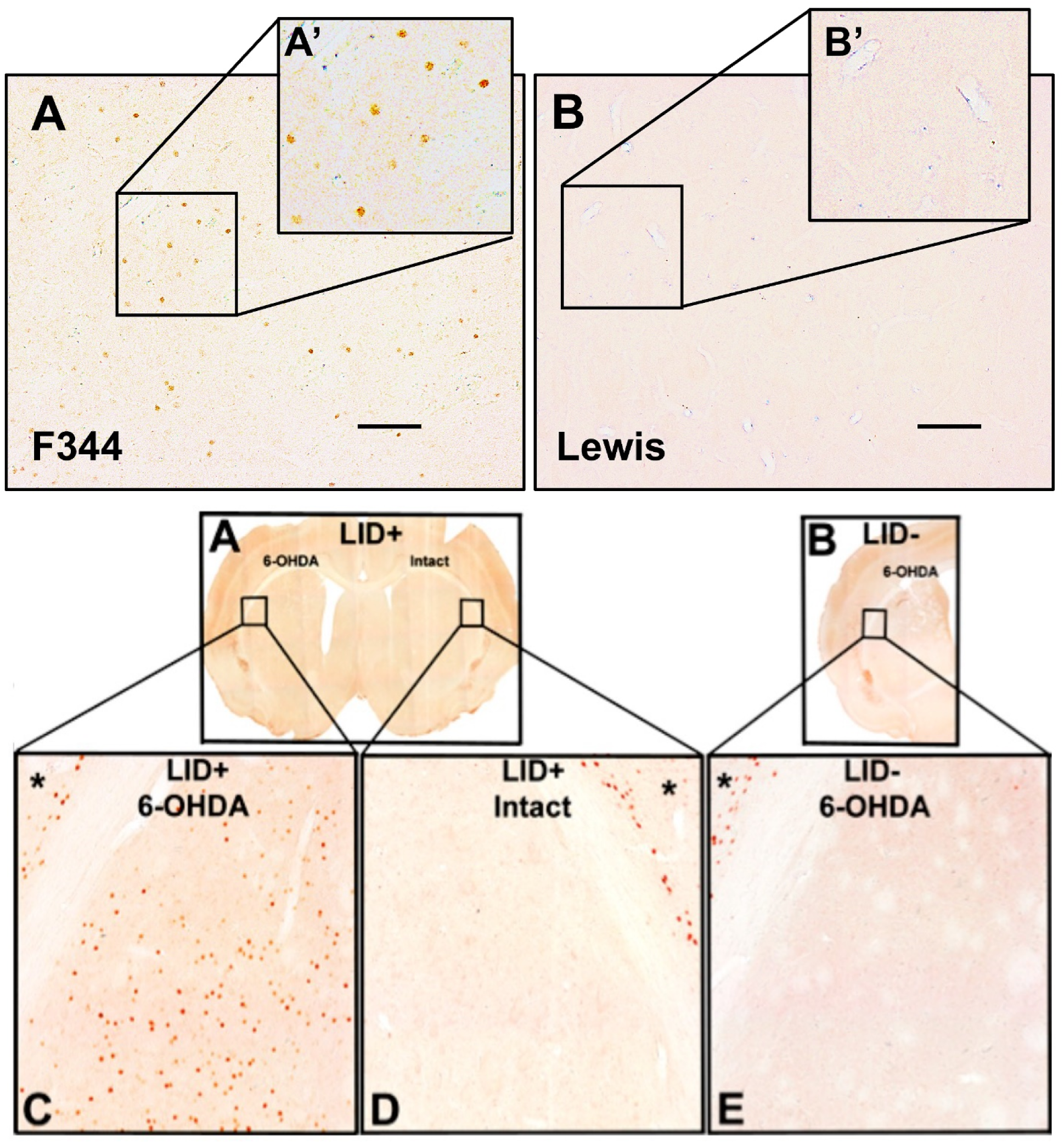
Abnormal Nurr1 upregulation in dyskinetic rats. (A) Nurr1 IHC in the lesioned striatum of a LID+ F344 rat. Abnormal Nurr1 expression can be seen throughout the lesioned striatum. (B) Nurr1 IHC in the lesioned striatum of an LID+ Lewis rat. No Nurr1 expression is seen in these animals. Scale bar=100µm.

**Supplemental Figure 2.**
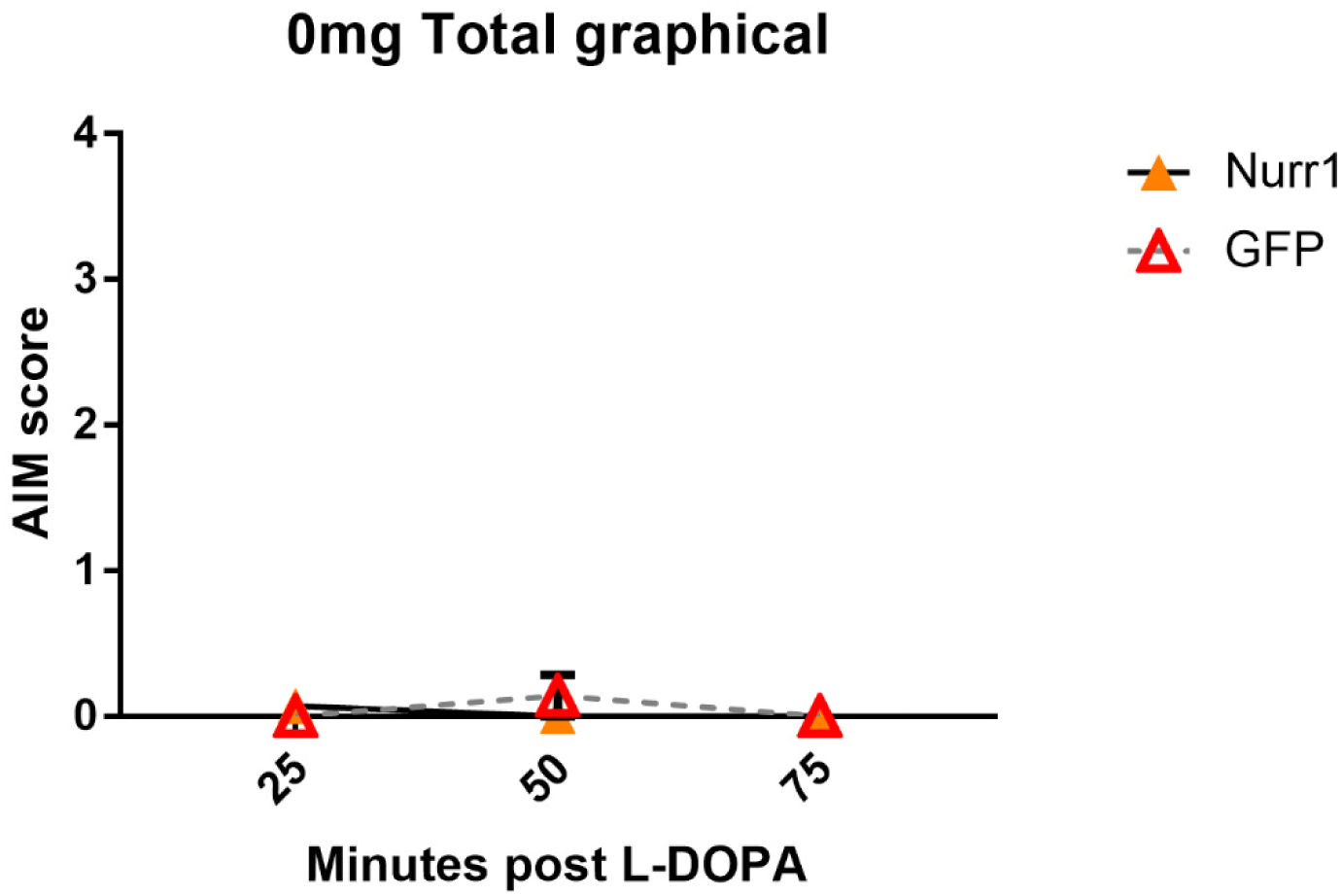
Striatal Nurr1 overexpression does not induce drug-independent AIMs. F344 rats treated with rAAV-Nurr1 or rAAV-GFP do not express AIMs when treated with vehicle (0mg/kg L-DOPA, 12mg/kg benserazide) showing that Nurr1 overexpression does not promote AIMs without drug treatment.

**Supplemental Figure 3.**
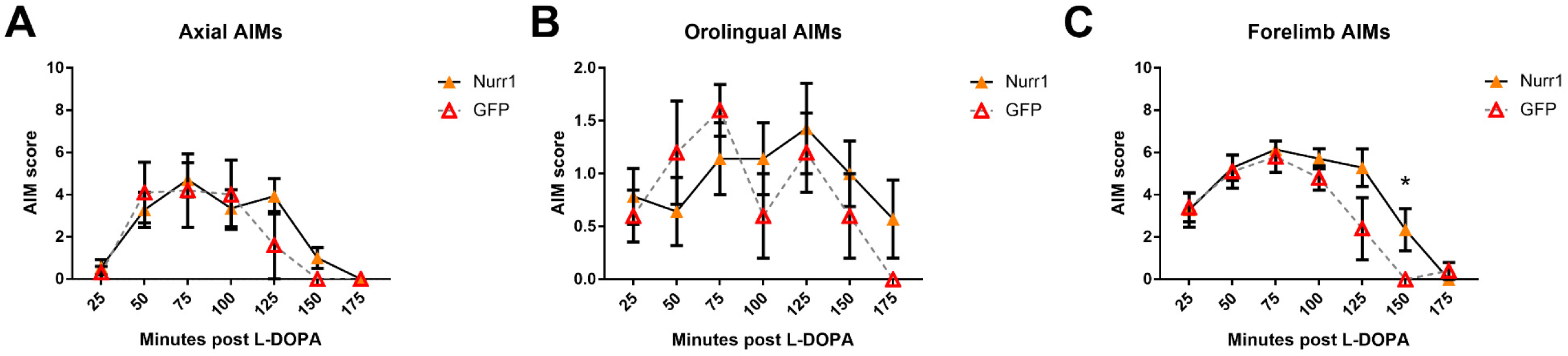
Nurr1 does not impact individual AOLs in LID-susceptible rats. (A) Axial AIMs—comprised of trunk and neck dystonia—are not different between rAAV-Nurr1 and rAAV-GFP F344 rats. (B) Orolingual AIMs—comprised of chewing and tongue protrusions—are not different between vector groups. (C) Forelimb AIMs—comprised of forelimb hyperkinesia and dystonia—are not different at most time points between vector treatment groups. There was a significant difference between the groups at 150 minutes post injection(*=p≤0.05). Rating period shown at day 17 with 8mg/kg L-DOPA dosing.

**Supplemental Figure 4.**
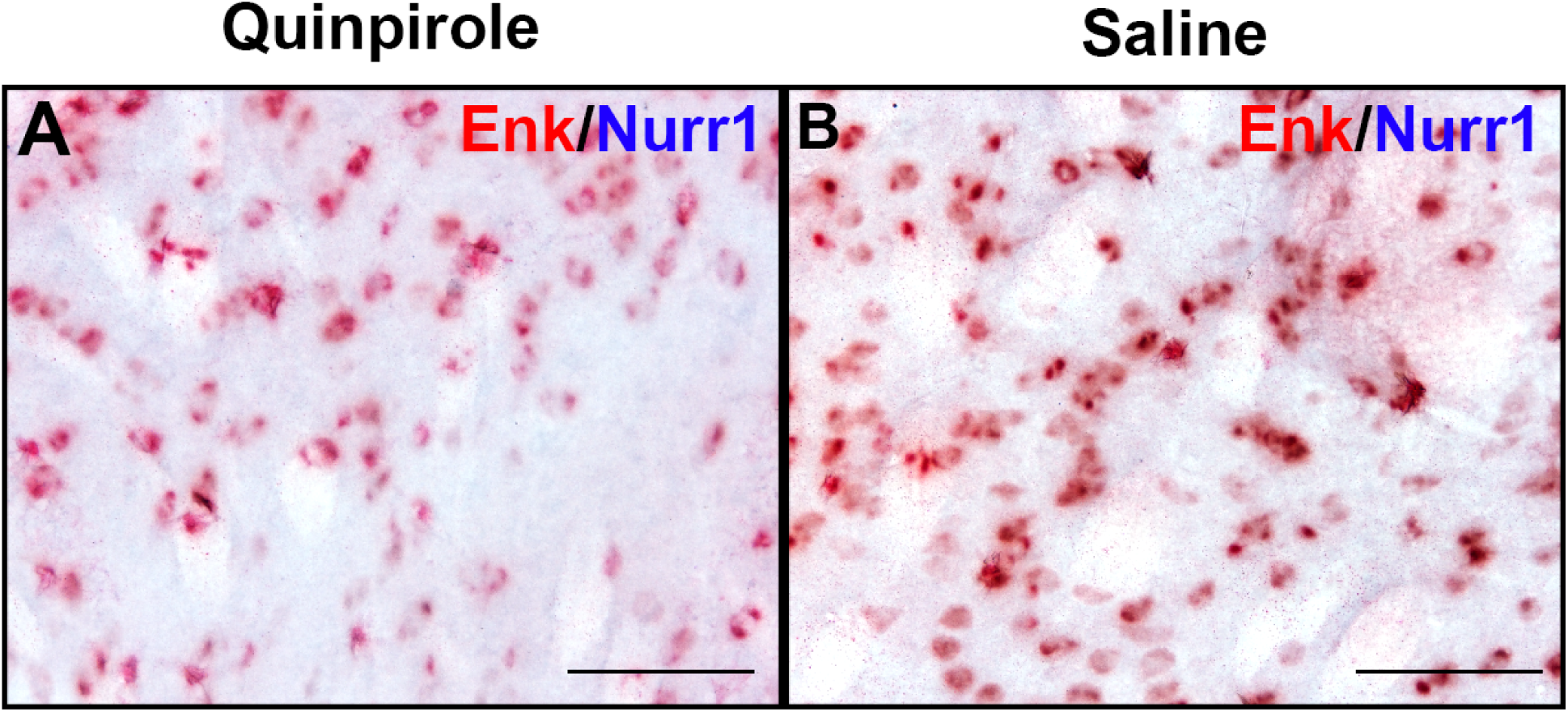
Nurr1 colocalization with iMSN in quinpirole and saline treated rats. *(A) In situ* hybridization for enkephalin (Enk) and with IHC for Nurr1 in a dyskinetic rat treated with quinpirole. No Nurr1 protein was observed in these animals. (B) *In situ* hybridization for Enk with IHC for Nurr1 in a saline treated animal show no abnormal Nurr1 induction in saline treated rats.

**Supplemental Figure 5.**
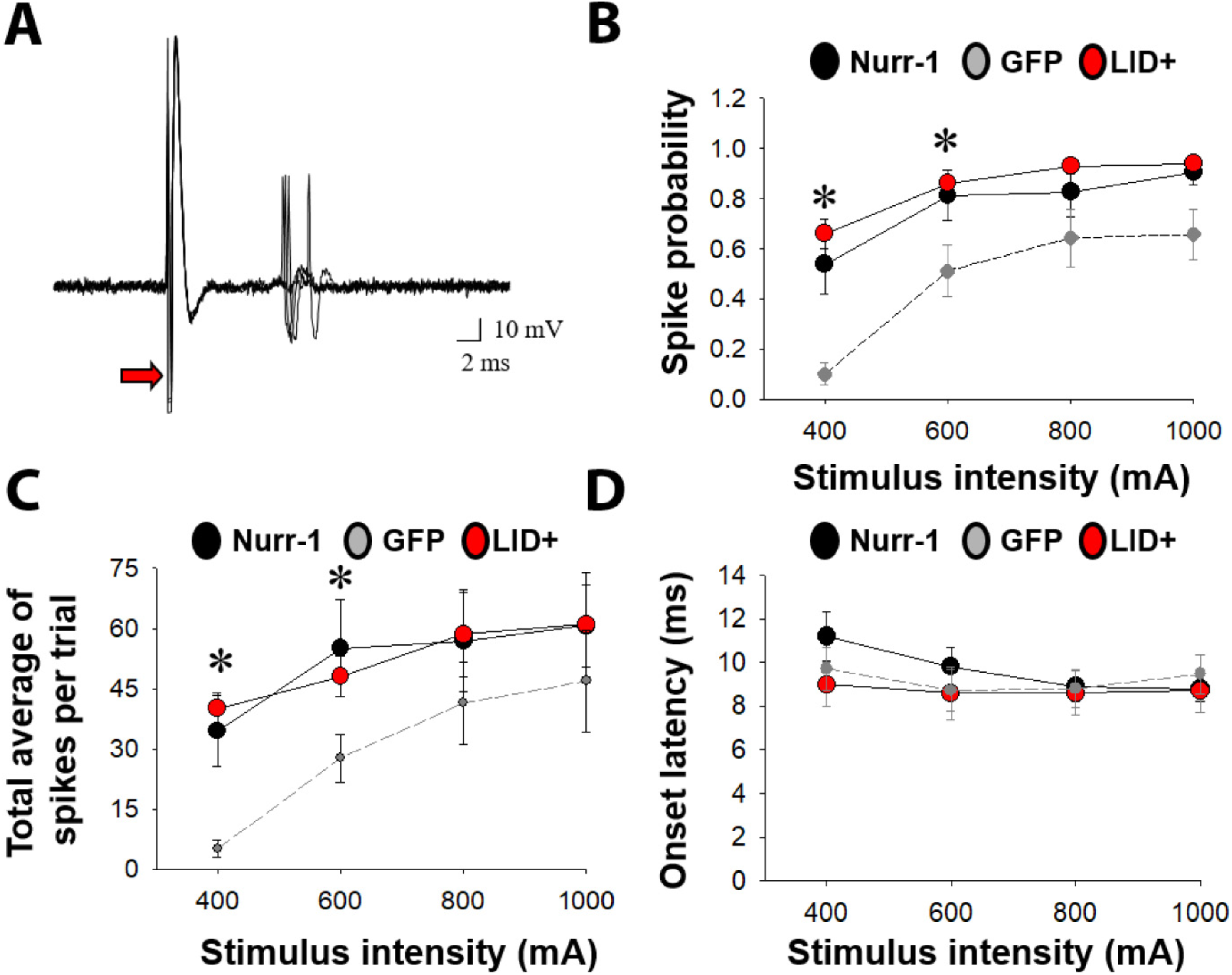
Cortically-evoked response of striatonigral MSN to antidromic stimulation. Comparisons between cortically-evoked spike characteristics of antidromically-activated striatonigral projection neurons recorded from DA-depleted rAAV-naïve rats treated with chronic L-DOPA (LID+), rAAV-GFP injected L-DOPA naïve animals, or rAAV-Nurr1 injected L-DOPA naïve animals. (A) Traces of typical cortically-evoked responses from isolated striatonigral projection neurons. Ten consecutive overlaid responses are shown. (B-D) graphs compare the spike probability (B), average number of total spikes evoked during a stimulus trial at each current intensity tested (C), and onset latency of evoked spikes (D) during cortical stimulation. Stimulus intensity-dependent effects on cortically-evoked spike probability of striatonigral projection neurons were observed in both LID+ and Nurr-1 overexpressing rats as compared to vehicle treated DA-depleted GFP-expressing controls (* p< 0.001). Post-hoc comparisons revealed a significant increase in the probability and number of evoked responses to cortical stimulation at the 400-600 µA current intensities in both LID+ and Nurr-1 overexpressing rats as compared to vehicle treated DA-depleted, GFP-expressing control rats (*p< 0.05). No significant differences in onset latency (p> 0.05) or SD of latency (data not shown) of cortically-evoked responses were observed.

## References Cited

1. Cotzias GC, Van Woert MH, Schiffer LM (1967): Aromatic amino acids and modification of parkinsonism. The New England journal of medicine. 276:374–379.

2. Fox SH, Katzenschlager R, Lim SY, Ravina B, Seppi K, Coelho M, et al. (2011): The Movement Disorder Society Evidence-Based Medicine Review Update: Treatments for the motor symptoms of Parkinson’s disease. Mov Disord. 26 Suppl 3:S2–41.

3. Ahlskog JE, Muenter MD (2001): Frequency of levodopa-related dyskinesias and motor fluctuations as estimated from the cumulative literature. Mov Disord. 16:448–458.

4. Manson A, Stirpe P, Schrag A (2012): Levodopa-induced-dyskinesias clinical features, incidence, risk factors, management and impact on quality of life. J Parkinsons Dis. 2:189–198.

5. Heiman M, Heilbut A, Francardo V, Kulicke R, Fenster RJ, Kolaczyk ED, et al. (2014): Molecular adaptations of striatal spiny projection neurons during levodopa-induced dyskinesia. Proceedings of the National Academy of Sciences of the United States of America. 111:4578–4583.

6. Xiao Q, Castillo SO, Nikodem VM (1996): Distribution of messenger RNAs for the orphan nuclear receptors Nurr1 and Nur77 (NGFI-B) in adult rat brain using in situ hybridization. Neuroscience. 75:221–230.

7. Manfredsson F, Kanaan M, Lipton J, J. Collier T, E. Caryl S, Cole-Strauss A, et al. (2014): Ectopic Nurr1 in striatal neurons results in enhanced levodopa-induced dyskinesias in the 6-OHDA rat model of Parkinson’s disease.

8. Sellnow RC, Steece-Collier K, Kanaan NM, Sortwell CE, Collier TJ, Cole-Strauss A, et al. (2015): 709. rAAV-Mediated Regulation of Striatal Nurr1 Expression Alters Development and Severity of Levodopa-Induced Dyskinesias in the 6-OHDA Rat Model of Parkinson’s Disease. Molecular Therapy. 23:S282–S283.

9. Sodersten E, Feyder M, Lerdrup M, Gomes AL, Kryh H, Spigolon G, et al. (2014): Dopamine signaling leads to loss of Polycomb repression and aberrant gene activation in experimental parkinsonism. PLoS genetics. 10:e1004574.

10. Giguere V (1999): Orphan nuclear receptors: from gene to function. Endocrine reviews. 20:689–725.

11. Zetterstrom RH, Solomin L, Jansson L, Hoffer BJ, Olson L, Perlmann T (1997): Dopamine neuron agenesis in Nurr1-deficient mice. Science. 276:248–250.

12. Jiang C, Wan X, He Y, Pan T, Jankovic J, Le W (2005): Age-dependent dopaminergic dysfunction in Nurr1 knockout mice. Exp Neurol. 191:154–162.

13. Kadkhodaei B, Ito T, Joodmardi E, Mattsson B, Rouillard C, Carta M, et al. (2009): Nurr1 is required for maintenance of maturing and adult midbrain dopamine neurons. The Journal of neuroscience : the official journal of the Society for Neuroscience. 29:15923–15932.

14. Le WD, Xu P, Jankovic J, Jiang H, Appel SH, Smith RG, et al. (2003): Mutations in NR4A2 associated with familial Parkinson disease. Nature genetics. 33:85–89.

15. Dekker MC, Bonifati V, van Duijn CM (2003): Parkinson’s disease: piecing together a genetic jigsaw. Brain. 126:1722–1733.

16. Kim C-H, Han B-S, Moon J, Kim D-J, Shin J, Rajan S, et al. (2015): Nuclear receptor Nurr1 agonists enhance its dual functions and improve behavioral deficits in an animal model of Parkinson’s disease.

17. Smith GA, Rocha EM, Rooney T, Barneoud P, McLean JR, Beagan J, et al. (2015): A Nurr1 agonist causes neuroprotection in a Parkinson’s disease lesion model primed with the toll-like receptor 3 dsRNA inflammatory stimulant poly(I:C). PloS one. 10:e0121072–e0121072.

18. Dong J, Li S, Mo JL, Cai HB, Le WD (2016): Nurr1-Based Therapies for Parkinson’s Disease. CNS neuroscience & therapeutics. 22:351–359.

19. Prescott IA, Liu LD, Dostrovsky JO, Hodaie M, Lozano AM, Hutchison WD (2014): Lack of depotentiation at basal ganglia output neurons in PD patients with levodopa-induced dyskinesia. Neurobiol Dis. 71:24–33.

20. Picconi B, Centonze D, Hakansson K, Bernardi G, Greengard P, Fisone G, et al. (2003): Loss of bidirectional striatal synaptic plasticity in L-DOPA-induced dyskinesia. Nat Neurosci. 6:501–506.

21. Ryan MB, Bair-Marshall C, Nelson AB (2018): Aberrant Striatal Activity in Parkinsonism and Levodopa-Induced Dyskinesia. Cell Rep. 23:3438–3446.e3435.

22. Zhang Y, Meredith GE, Mendoza-Elias N, Rademacher DJ, Tseng KY, Steece-Collier K (2013): Aberrant restoration of spines and their synapses in L-DOPA-induced dyskinesia: involvement of corticostriatal but not thalamostriatal synapses. The Journal of neuroscience : the official journal of the Society for Neuroscience. 33:11655–11667.

23. Fieblinger T, Graves SM, Sebel LE, Alcacer C, Plotkin JL, Gertler TS, et al. (2014): Cell type-specific plasticity of striatal projection neurons in parkinsonism and L-DOPA-induced dyskinesia. Nature communications. 5:5316–5316.

24. Nishijima H, Suzuki S, Kon T, Funamizu Y, Ueno T, Haga R, et al. (2014): Morphologic changes of dendritic spines of striatal neurons in the levodopa-induced dyskinesia model. Movement Disorders. 29:336–343.

25. Suárez LM, Solís O, Caramés JM, Taravini IR, Solís JM, Murer MG, et al. (2014): L-DOPA treatment selectively restores spine density in dopamine receptor D2-expressing projection neurons in dyskinetic mice. Biological psychiatry. 75:711–722.

26. Maiti P, Manna J, Ilavazhagan G, Rossignol J, Dunbar GL (2015): Molecular regulation of dendritic spine dynamics and their potential impact on synaptic plasticity and neurological diseases. Neuroscience and biobehavioral reviews. 59:208–237.

27. Bagetta V, Ghiglieri V, Sgobio C, Calabresi P, Picconi B (2010): Synaptic dysfunction in Parkinson’s disease. Biochemical Society transactions. 38:493–497.

28. Spiga S, Mulas G, Piras F, Diana M (2014): The “addicted” spine. Frontiers in neuroanatomy. 8:110–110.

29. McNeill TH, Brown SA, Rafols JA, Shoulson I (1988): Atrophy of medium spiny I striatal dendrites in advanced Parkinson’s disease. Brain Res. 455:148–152.

30. Villalba RM, Smith Y (2018): Loss and remodeling of striatal dendritic spines in Parkinson’s disease: from homeostasis to maladaptive plasticity? J Neural Transm (Vienna). 125:431–447.

31. Suarez LM, Solis O, Aguado C, Lujan R, Moratalla R (2016): L-DOPA Oppositely Regulates Synaptic Strength and Spine Morphology in D1 and D2 Striatal Projection Neurons in Dyskinesia. Cereb Cortex. 26:4253–4264.

32. Peña de Ortiz S, Maldonado-Vlaar CS, Carrasquillo Y (2000): Hippocampal expression of the orphan nuclear receptor gene hzf-3/nurr1 during spatial discrimination learning. Neurobiology of learning and memory. 74:161–178.

33. Hawk JD, Bookout AL, Poplawski SG, Bridi M, Rao AJ, Sulewski ME, et al. (2012): NR4A nuclear receptors support memory enhancement by histone deacetylase inhibitors. Journal of Clinical Investigation. 122:3593–3602.

34. Colon-Cesario WI, Martinez-Montemayor MM, Morales S, Felix J, Cruz J, Adorno M, et al. (2006): Knockdown of Nurr1 in the rat hippocampus: implications to spatial discrimination learning and memory. Learning & memory (Cold Spring Harbor, NY). 13:734–744.

35. Campos-Melo D, Galleguillos D, Sanchez N, Gysling K, Andres ME (2013): Nur transcription factors in stress and addiction. Frontiers in molecular neuroscience. 6:44.

36. Benskey MJ, Sandoval IM, Manfredsson FP (2016): Continuous Collection of Adeno-Associated Virus from Producer Cell Medium Significantly Increases Total Viral Yield. Human gene therapy methods. 27:32–45.

37. Sandoval IM, Kuhn NM, Manfredsson FP (2019): Multimodal Production of Adeno-Associated Virus. Methods Mol Biol. 1937:101–124.

38. Benskey MJ, Manfredsson FP (2016): Intraparenchymal Stereotaxic Delivery of rAAV and Special Considerations in Vector Handling. Methods Mol Biol. 1382:199–215.

39. Sellnow RC, Newman JH, Chambers N, West AR, Steece-Collier K, Sandoval IM, et al. (2019): Regulation of dopamine neurotransmission from serotonergic neurons by ectopic expression of the dopamine D2 autoreceptor blocks levodopa-induced dyskinesia. Acta neuropathologica communications. 7:8.

40. Manfredsson FP, Burger C, Sullivan LF, Muzyczka N, Lewin AS, Mandel RJ (2007): rAAV-mediated nigral human parkin over-expression partially ameliorates motor deficits via enhanced dopamine neurotransmission in a rat model of Parkinson’s disease. Exp Neurol. 207:289–301.

41. Schallert T (2006): Behavioral tests for preclinical intervention assessment. NeuroRX. 3:497–504.

42. Lundblad M, Andersson M, Winkler C, Kirik D, Wierup N, Cenci MA (2002): Pharmacological validation of behavioural measures of akinesia and dyskinesia in a rat model of Parkinson’s disease. The European journal of neuroscience. 15:120–132.

43. Steece-Collier K, Collier TJ, Danielson PD, Kurlan R, Yurek DM, Sladek JR (2003): Embryonic mesencephalic grafts increase levodopa-induced forelimb hyperkinesia in parkinsonian rats. Movement disorders : official journal of the Movement Disorder Society. 18:1442–1454.

44. Maries E, Kordower JH, Chu Y, Collier TJ, Sortwell CE, Olaru E, et al. (2006): Focal not widespread grafts induce novel dyskinetic behavior in parkinsonian rats. Neurobiology of disease. 21:165–180.

45. Simuni T, Siderowf A, Lasch S, Coffey CS, Caspell-Garcia C, Jennings D, et al. (2018): Longitudinal Change of Clinical and Biological Measures in Early Parkinson’s Disease: Parkinson’s Progression Markers Initiative Cohort. Mov Disord. 33:771–782.

46. Zesiewicz TA, Chriscoe S, Jimenez T, Upward J, Davy M, VanMeter S (2017): A randomized, fixed-dose, dose-response study of ropinirole prolonged release in advanced Parkinson’s disease. Neurodegenerative disease management. 7:61–72.

47. Zesiewicz TA, Chriscoe S, Jimenez T, Upward J, VanMeter S (2017): A fixed-dose, dose-response study of ropinirole prolonged release in early stage Parkinson’s disease. Neurodegenerative disease management. 7:49–59.

48. Hauser RA, Rascol O, Korczyn AD, Jon Stoessl A, Watts RL, Poewe W, et al. (2007): Ten-year follow-up of Parkinson’s disease patients randomized to initial therapy with ropinirole or levodopa. Mov Disord. 22:2409–2417.

49. Bastide MF, Meissner WG, Picconi B, Fasano S, Fernagut P-O, Feyder M, et al. (2015): Pathophysiology of L-dopa-induced motor and non-motor complications in Parkinson’s disease. Progress in neurobiology. 132:96–168.

50. Lundblad M, Usiello A, Carta M, Hakansson K, Fisone G, Cenci MA (2005): Pharmacological validation of a mouse model of l-DOPA-induced dyskinesia. Exp Neurol. 194:66–75.

51. Carta M, Carlsson T, Munoz A, Kirik D, Bjorklund A (2008): Involvement of the serotonin system in L-dopa-induced dyskinesias. Parkinsonism & related disorders. 14 Suppl 2:S154–158.

52. Lane EL, Dunnett SB (2010): Pre-treatment with dopamine agonists influence L-dopa mediated rotations without affecting abnormal involuntary movements in the 6-OHDA lesioned rat. Behavioural brain research. 213:66–72.

53. Threlfell S, Sammut S, Menniti FS, Schmidt CJ, West AR (2009): Inhibition of Phosphodiesterase 10A Increases the Responsiveness of Striatal Projection Neurons to Cortical Stimulation. The Journal of pharmacology and experimental therapeutics. 328:785–795.

54. Sammut S, Threlfell S, West AR (2010): Nitric oxide-soluble guanylyl cyclase signaling regulates corticostriatal transmission and short-term synaptic plasticity of striatal projection neurons recorded in vivo. Neuropharmacology. 58:624–631.

55. Lu XY, Ghasemzadeh MB, Kalivas PW (1998): Expression of D1 receptor, D2 receptor, substance P and enkephalin messenger RNAs in the neurons projecting from the nucleus accumbens. Neuroscience. 82:767–780.

56. Levine ND, Rademacher DJ, Collier TJ, O’Malley JA, Kells AP, San Sebastian W, et al. (2013): Advances in thin tissue Golgi-Cox impregnation: Fast, reliable methods for multi-assay analyses in rodent and non-human primate brain. Journal of Neuroscience Methods. 213:214–227.

57. Polinski NK, Gombash SE, Manfredsson FP, Lipton JW, Kemp CJ, Cole-Strauss A, et al. (2015): Recombinant adenoassociated virus 2/5-mediated gene transfer is reduced in the aged rat midbrain. Neurobiology of aging. 36:1110–1120.

58. Padovan-Neto FE, Sammut S, Chakroborty S, Dec AM, Threlfell S, Campbell PW, et al. (2015): Facilitation of corticostriatal transmission following pharmacological inhibition of striatal phosphodiesterase 10A: role of nitric oxide-soluble guanylyl cyclase-cGMP signaling pathways. J Neurosci. 35:5781–5791.

59. Schwarting RK, Huston JP (1996): The unilateral 6-hydroxydopamine lesion model in behavioral brain research. Analysis of functional deficits, recovery and treatments. Prog Neurobiol. 50:275–331.

60. Soderstrom KE, O’Malley JA, Levine ND, Sortwell CE, Collier TJ, Steece-Collier K (2010): Impact of dendritic spine preservation in medium spiny neurons on dopamine graft efficacy and the expression of dyskinesias in parkinsonian rats. The European journal of neuroscience. 31:478–490.

61. Fieblinger T, Cenci MA (2015): Zooming in on the small: the plasticity of striatal dendritic spines in L-DOPA-induced dyskinesia. Movement disorders : official journal of the Movement Disorder Society. 30:484–493.

62. Brodkin ES, Carlezon WA, Jr., Haile CN, Kosten TA, Heninger GR, Nestler EJ (1998): Genetic analysis of behavioral, neuroendocrine, and biochemical parameters in inbred rodents: initial studies in Lewis and Fischer 344 rats and in A/J and C57BL/6J mice. Brain Res. 805:55–68.

63. Fole A, Miguens M, Morales L, Gonzalez-Martin C, Ambrosio E, Del Olmo N (2017): Lewis and Fischer 344 rats as a model for genetic differences in spatial learning and memory: Cocaine effects. Progress in neuro-psychopharmacology & biological psychiatry. 76:49–57.

64. Miguens M, Coria SM, Higuera-Matas A, Fole A, Ambrosio E, Del Olmo N (2011): Depotentiation of hippocampal long-term potentiation depends on genetic background and is modulated by cocaine self-administration. Neuroscience. 187:36–42.

65. Valenza M, Picetti R, Yuferov V, Butelman ER, Kreek MJ (2016): Strain and cocaine-induced differential opioid gene expression may predispose Lewis but not Fischer rats to escalate cocaine self-administration. Neuropharmacology. 105:639–650.

66. Maira M, Martens C, Philips A, Drouin J (1999): Heterodimerization between members of the Nur subfamily of orphan nuclear receptors as a novel mechanism for gene activation. Molecular and cellular biology. 19:7549–7557.

67. Mahmoudi S, Samadi P, Gilbert F, Ouattara B, Morissette M, Grégoire L, et al. (2009): Nur77 mRNA levels and L-Dopa-induced dyskinesias in MPTP monkeys treated with docosahexaenoic acid. Neurobiology of disease. 36:213–222.

68. St-Hilaire M, Landry E, Levesque D, Rouillard C (2003): Denervation and repeated L-DOPA induce a coordinate expression of the transcription factor NGFI-B in striatal projection pathways in hemi-parkinsonian rats. Neurobiol Dis. 14:98–109.

69. Rouillard C, Baillargeon J, Paquet B, St-Hilaire M, Maheux J, Levesque C, et al. (2018): Genetic disruption of the nuclear receptor Nur77 (Nr4a1) in rat reduces dopamine cell loss and l-Dopa-induced dyskinesia in experimental Parkinson’s disease. Exp Neurol. 304:143–153.

70. Chen Y, Wang Y, Ertürk A, Kallop D, Jiang Z, Weimer RM, et al. (2014): Activity-induced Nr4a1 regulates spine density and distribution pattern of excitatory synapses in pyramidal neurons. Neuron. 83:431–443.

71. Yamada H, Kuroki T, Nakahara T, Hashimoto K, Tsutsumi T, Hirano M, et al. (2007): The dopamine D1 receptor agonist, but not the D2 receptor agonist, induces gene expression of Homer 1a in rat striatum and nucleus accumbens. Brain Res. 1131:88–96.

72. Pollack AE, Yates TM (1999): Prior D1 dopamine receptor stimulation is required to prime D2-mediated striatal Fos expression in 6-hydroxydopamine-lesioned rats. Neuroscience. 94:505–514.

73. Pollack AE, Thomas LI (2010): D1 priming enhances both D1- and D2-mediated rotational behavior and striatal Fos expression in 6-hydroxydopamine lesioned rats. Pharmacology, biochemistry, and behavior. 94:346–351.

74. Decressac M, Volakakis N, Björklund A, Perlmann T (2013): NURR1 in Parkinson disease—from pathogenesis to therapeutic potential. Nature Reviews Neurology. 9:629–636.

75. De Miranda BR, Popichak KA, Hammond SL, Jorgensen BA, Phillips AT, Safe S, et al. (2015): The Nurr1 Activator 1,1-Bis(3’-Indolyl)-1-(p-Chlorophenyl)Methane Blocks Inflammatory Gene Expression in BV-2 Microglial Cells by Inhibiting Nuclear Factor kappaB. Molecular pharmacology. 87:1021–1034.

76. Volakakis N, Tiklova K, Decressac M, Papathanou M, Mattsson B, Gillberg L, et al. (2015): Nurr1 and Retinoid X Receptor Ligands Stimulate Ret Signaling in Dopamine Neurons and Can Alleviate α-Synuclein Disrupted Gene Expression. The Journal of neuroscience : the official journal of the Society for Neuroscience. 35:14370–14385.

77. Zetterstrom RH, Williams R, Perlmann T, Olson L (1996): Cellular expression of the immediate early transcription factors Nurr1 and NGFI-B suggests a gene regulatory role in several brain regions including the nigrostriatal dopamine system. Brain research Molecular brain research. 41:111–120.

